# RNA isoform-resolved multiplexed sequencing with bioorthogonal barcoding

**DOI:** 10.64898/2026.09.22.753383

**Authors:** Turnee N. Malik, Joshua Berlind, Amina Khan, Bryan van Nimwegen, Mislav Acman, Neal D. Amin

**Affiliations:** Sculpta, Inc., 135 Mississippi St, San Francisco CA, 94017, USA; Bryan Bioinformatics, Wageningen, The Netherlands; Omics d.o.o., Manterovcak 72, 10000, Zagreb, Croatia

## Abstract

RNA isoform dysregulation drives disease pathogenesis and is the target of FDA-approved splice-switching therapeutics. However, multiplexed sequencing methods discard splice junction information because only 3’ termini are barcoded and counted. Here, we repurpose acylation and click chemistries to conjugate bioorthogonal barcodes (bobcodes) onto multiple internal positions along cellular RNAs. Bobcoded RNAs from multiple samples are pooled for multiplexed cDNA synthesis, during which reverse transcriptase switches from each RNA template onto its tethered bobcode with greater than 99% accuracy in species mixing experiments. Bobcode attachment intervals set cDNA insert sizes without a library fragmentation step, and priming with poly(dT) or random hexamers selects between 3’-end counting and full-length isoform capture. A bioorthogonal barcode-sequencing (BOB-seq v0.1) drug screen identifies transcriptome-wide on- and off-target RNA splicing effects and outperforms existing multiplexing RNA sequencing methods in workflow simplicity, sample-to-sample variability, and barcoding accuracy. Bobcodes add isoform resolution to scalable multiplexed RNA sequencing.

## Introduction

Dysregulation of RNA splicing alters isoform diversity and drives a wide range of human conditions including neurodegeneration, cancer, cardiomyopathy, and aging^1–4^. Modifying RNA splicing is an established therapeutic mechanism of action. The FDA-approved antisense oligonucleotide (ASO) drug nusinersen redirects SMN2 splice site selection to restore protein production in spinal muscular atrophy (SMA), reducing ventilator need and death^5^. Splice-switching ASOs have also demonstrated disease-modifying potential in clinical trials for Dravet syndrome, reducing seizure frequency in patients already on conventional anti-epileptics^6^. The FDA approval of risdiplam for SMA has demonstrated that small molecules are an alternative and orally bioavailable modality to module RNA splicing in patients^7^.

To accelerate the development of splice-switching drugs, ASO and small-molecule libraries must be screened at high throughput for on- and off-target effects on RNA isoforms. Existing multiplexed transcriptomic methods, including DRUG-seq and prime-seq, capture only the 3’ ends of transcripts and were built to profile drug effects on gene expression rather than transcriptome-wide splicing^8–11^. In these methods, sample-specific barcodes are incorporated into cDNA during reverse transcription, and the barcoded libraries are then pooled for multiplexed processing. Reducing library size for short-read sequencing requires fragmentation or tagmentation, which separates splicing information in the gene body from the poly(dT) priming barcodes confined to the 3’ termini.

Random priming along the gene body is not a practical alternative, because rRNA dominates unless an RNA selection step precedes reverse transcription on each individual sample. Adapting existing enzymatic barcoding methods for isoform capture would instead require long-read sequencing, which yields two orders of magnitude fewer reads per run, adds several time-consuming library preparation steps, and therefore raises the cost per sample substantially at comparable sequencing depth^12,13^. Because of these limitations of barcoding with a reverse transcription primer, no highly multiplexed, isoform-resolved platform has been described for scalable drug screens.

We reasoned that reverse transcription-based barcoding platforms are intrinsically limited in their ability to capture RNA isoforms because barcodes are placed only at the ends of cDNAs, far from most RNA splice junctions. Instead, we considered whether barcodes could be conjugated along the length of transcripts where RNA splicing information resides, rather than just the termini of cDNA.

Two chemistries suggested a route (**Fig. 1A**): (1) cell-permeable RNA structure probing agents install azide-bearing covalent adducts on internal positions of the RNA backbone, chemically decorating native RNAs in living cells within minutes^14–19^; and (2) bioorthogonal click chemistry chemoselectively couples azides with strained alkynes in complex biological environments, without catalysts or enzymes^20–24^. Directly conjugating barcodes to RNA would enable early multiplexing, prior to enzymatic processing steps including reverse transcription (**Fig. 1B**).

**Figure 1.**
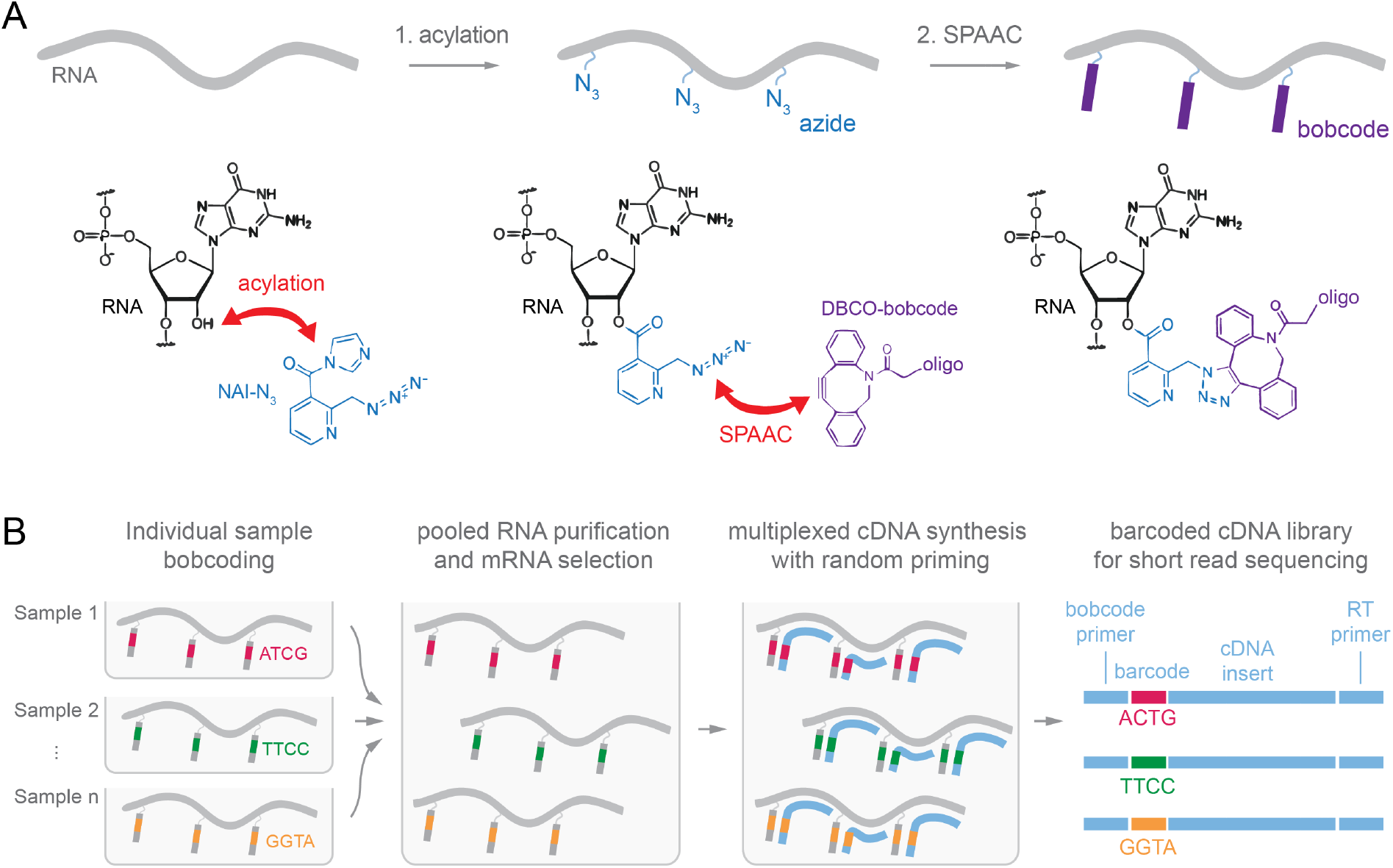
Bioorthogonal barcoding. (A) 2-(Azidomethyl)nicotinic acid imidazolide (NAI-N₃) acylates 2′-hydroxyl groups of native RNA, installing azide groups. An oligonucleotide carrying a dibenzocyclooctyne (DBCO) group then reacts with azide by strain-promoted azide–alkyne cycloaddition (SPAAC), covalently attaching the oligonucleotide to the RNA via click chemistry. (B) The RNA of each sample is conjugated to a bioorthogonal barcode (bobcode) bearing a unique barcode sequence. Samples are then pooled and reverse transcribed, producing a barcoded cDNA library for short-read sequencing capturing full length transcript information, without a fragmentation step.

Here, we introduce bioorthogonal barcodes (“bobcodes”), oligonucleotide barcodes attached at multiple internal positions along RNA, enzyme-free by two-step acylation and bioorthogonal click chemistries. Bobcoded RNAs are then pooled and purified for multiplexed cDNA synthesis, in which reverse transcriptases engage in intramolecular template switching from the RNA insert to the conjugated bobcode with greater than 99% accuracy. In a 24-sample multiplexed RNA sequencing experiment, we demonstrate that bioorthogonal barcode-sequencing (BOB-seq v0.1) captures transcriptome-wide small molecule and ASO drug effects on RNA isoforms missed by 3’ end barcoding methods. Benchmarked against DRUG-seq, BOB-seq achieved ∼10-fold less barcode swapping rates, shorter bench workflow duration and fewer steps, ∼25-fold higher capture of RNA splicing events, with lower inter-replicate variability in gene expression, demonstrating the effectiveness of chemical barcoding to improve transcriptomic data quality without compromising scale.

## Results

### RNA acylation efficiency and effects on cDNA synthesis

2-(azidomethyl)nicotinic acid imidazolide (NAI-N₃) is a membrane-permeable acylating reagent that installs an acyl adduct carrying an azide group onto the reactive 2′-hydroxyl position of RNA^14–16,19^. We developed a quantitative fluorescence-based assay to determine the extent of azide group installation on RNAs. First, we incubated *in vitro* total RNA with different concentrations of NAI-N₃ for 10 minutes at 37°C, purified RNA from excess NAI-N₃. We then added 1mM dibenzocyclooctyne-carboxyfluorescein (DBCO-FAM) to fluorescently label azide groups on RNA to saturation in a catalyst and enzyme-free strain-promoted azide-alkyne cycloaddition (SPAAC) click conjugation reaction (**Fig. 2A**). After 30 minutes, we purified RNA from excess DBCO-FAM. Fluorescence was quantified by spectrophotometer and RNA weight by nanodrop, providing a quantitative calculation of per-nucleotide azide labeling rates. RNAs exposed to higher concentrations of NAI-N_3_ exhibited higher fluorescence per nucleotide, and exposure to over 100mM NAI-N_3_ resulted in a calculated labeling rate of approximately one azide per 300 nucleotides (**Fig. 2B**).

**Fig 2.**
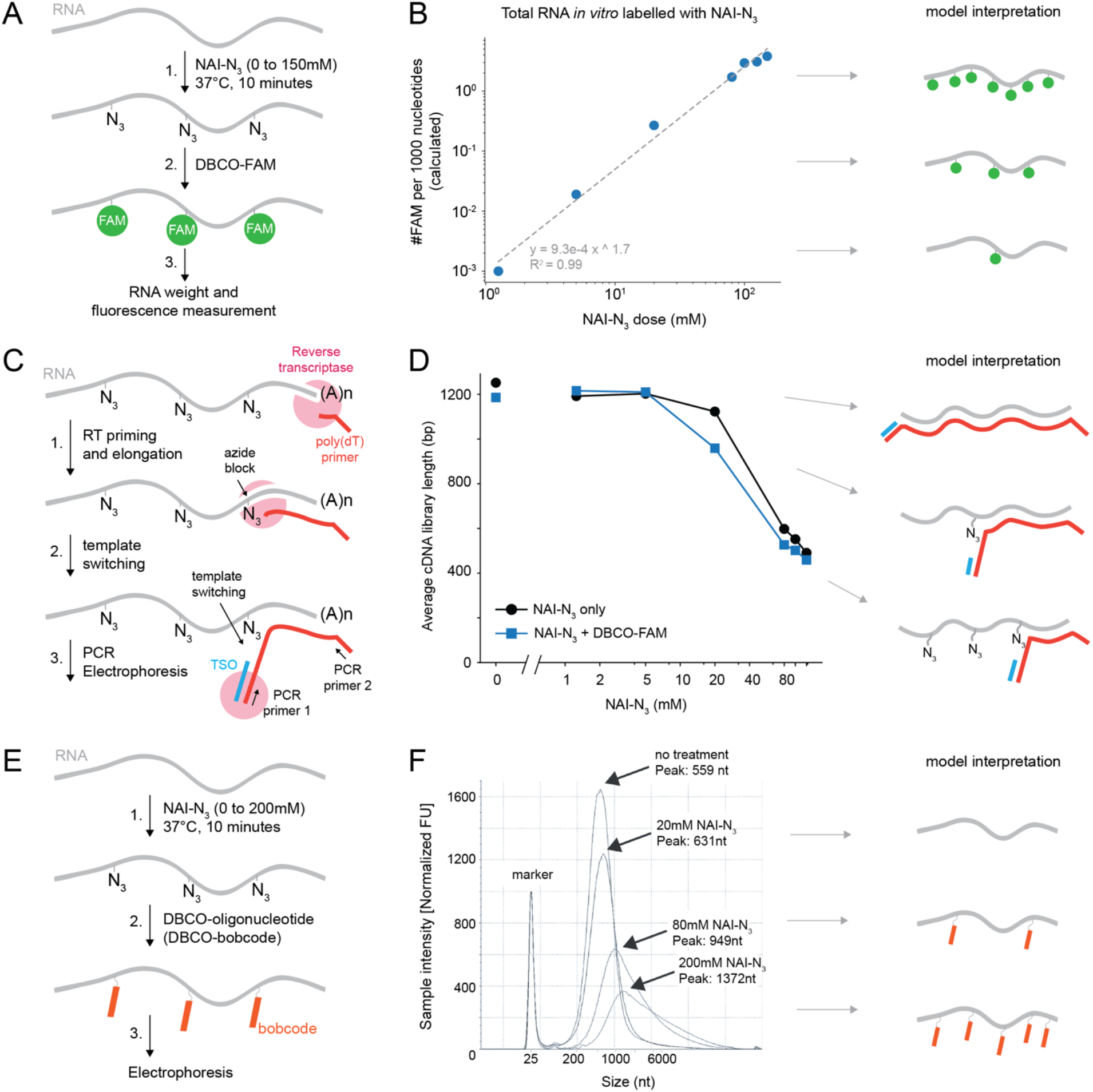
Acylation of RNA tunes cDNA library size and extent of oligonucleotide click conjugation to RNA. (A) RNA is acylated with NAI-N_3_ then labeled with DBCO-FAM by SPAAC chemistry. (B) FAM incorporation per 1000 nucleotides increases with NAI-N_3_ concentration (log-log). (C) Poly(dT)-primed reverse transcription of NAI-N_3_-labeled RNA. Internal adducts stall the reverse transcriptase and trigger template switching onto a template-switching oligonucleotide (TSO). (D) Average cDNA library length (adapters plus insert) decreases with increased NAI-N_3_ concentration. (E) Fragmented RNA is acylated with NAI-N_3_, purified, and conjugated to 50-nt DBCO-oligonucleotides by SPAAC click chemistry. (F) Electrophoresis results of purified conjugates demonstrate higher NAI-N_3_ exposure increases the conjugation rate of bobcodes to RNA.

2′-OH acylation has been reported to protect RNA from both thermal and enzymatic degradation^25,26^. We assessed if acylation chemistry affects RNA stability by treating *in vitro* purified RNA or RNA within living cells with different concentrations of NAI-N_3_ for different durations. After RNA isolation and electrophoresis measurements, we observed that RINe scores were unaffected (**Fig. S1A,B**).

We next assessed how covalent backbone adducts impact cDNA library preparation steps including reverse transcriptase template switching. RNA adducts are known to stall or halt reverse transcriptases^27–31^, and RT pausing can promote strand transfer between RNA templates^32–35^ but, to the best of our knowledge, have not been applied in general cDNA preparation workflows. We performed reverse transcription with a poly(dT) primer and a template switching oligonucleotide (TSO) using total RNA treated with a range of NAI-N₃ concentrations from 0 to 125mM at 37°C for 10 minutes. DBCO-FAM was also added to azide-labeled RNA to assess the effect of a larger molecular weight adduct on reverse transcription. We then measured average cDNA library size with electrophoresis measurements after PCR amplification using primers for the reverse transcription primer and the TSO (**Fig. 2C**). cDNA library size decreased in a dose-dependent manner with higher NAI-N₃ concentrations, with or without subsequent conjugation with DBCO-FAM (**Fig. 2D**). These results demonstrate that 2’-O-acyl adducts terminate reverse transcription and trigger template switching at internal positions on RNA, and that cDNA library size can be dose-dependently tuned by NAI-N₃ treatment of RNA.

### Covalently conjugating DBCO-oligonucleotides to azide-labeled RNA with bioorthogonal chemistry

Next, we determined whether oligonucleotides conjugated to a DBCO could be chemically conjugated to azide-labeled RNA in a SPAAC click conjugation reaction. We fragmented total RNA with magnesium and heat to a standard size distribution, purified it, then treated it with a range of NAI-N₃ concentrations for 10 min at 37°C. After removing excess NAI-N₃, we incubated the azide-labeled RNA with a DBCO-labeled 50-mer DNA oligonucleotide (**Fig. 2E**). High conjugation efficiency required high NaCl and PEG in the SPAAC reaction buffer, conditions that screen the electrostatic repulsion between the two polyanions and raise their effective local concentration through macromolecular crowding (data not shown). We carried out SPAAC in 1.0X SPRI beads for 30 min at 37°C, followed by two rounds of SPRI cleanup to remove unconjugated DBCO-labeled oligonucleotides. Electrophoresis demonstrated that RNA pre-treated with higher concentrations of NAI-N₃ exhibited slower RNA migration (**Fig. 2F**), consistent with dose-dependent conjugation of DBCO-oligonucleotides and the resulting increase in molecular size of the RNA-oligonucleotide conjugates.

### Mechanisms to transfer bioorthogonal barcode (bobcode) sequence information to cDNA libraries

Bioorthogonal barcodes (“bobcodes”) covalently attached at 2’-OH positions are oriented orthogonal to the phosphodiester backbone of RNA, so sequence information cannot be directly captured by a reverse transcriptase synthesizing cDNA 5’ to 3’ along the RNA template. Capturing bobcode and RNA sequence information within cDNA libraries therefore required the development of a bobcode transfer mechanism.

We designed three transfer mechanisms (**Fig. 3A**), each leveraging the physical proximity of a bobcode to its own RNA or RNA:cDNA hybrid: (A) splint-mediated ligation of bobcodes to the reverse transcription primers incorporated into hybridized cDNA; (B) a poly(dT)-VN sequence at the 3’ end of bobcodes to enable intramolecular reverse transcription priming; and (C) template switching from the RNA onto the 3’ end of the bobcode during reverse transcription, with either the bobcode’s 3’ end or its 5’ end attached to DBCO.

**Figure 3.**
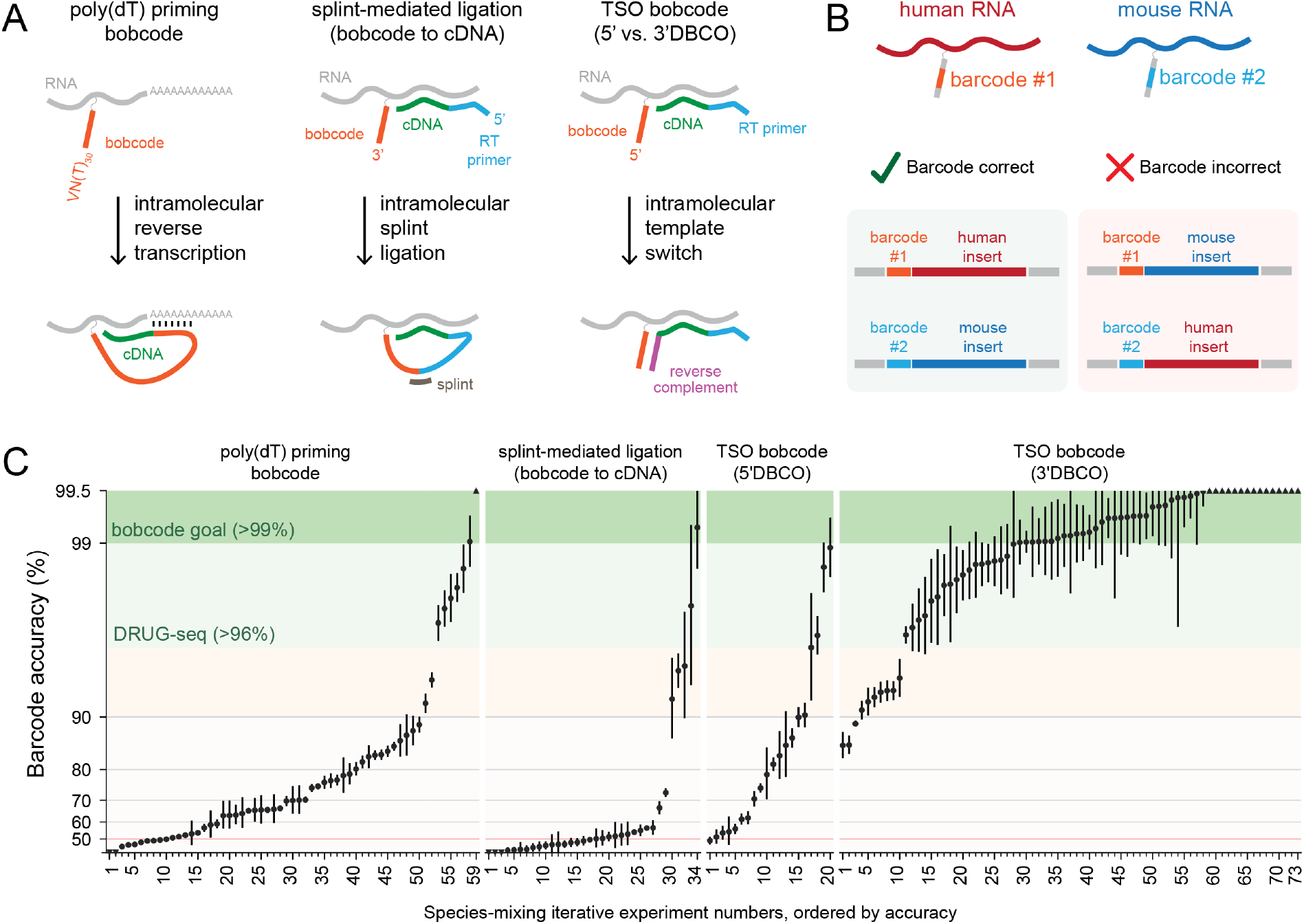
Greater than 99.5% barcoding accuracy using bioorthogonal barcode (bobcode) transfer mechanisms using reverse transcriptase template switching. (**A**) Three transfer mechanisms to connect bobcodes with cDNA hybridized to RNA templates. (**B**) Matching of barcodes with insert alignments to mouse or human transcriptomes to determine barcoding accuracy. (**C**) Barcode accuracy rates determined from independent sequencing experiments after iterative optimizations, arranged from low to high within each transfer mechanism category. Each data point represents an individual sequencing experiment. 95% confidence intervals shown reflecting only counting error from each sequencing experiment (Wilson score interval). Conditions described in (**Supplementary Table S1**).

Bobcoded RNAs pooled from multiple samples share a single reaction vessel, so transfer mechanisms are at risk of barcode swapping if the transfer occurs intermolecularly instead of intramolecularly. Therefore, we evaluated barcoding accuracy from each transfer mechanism in species-mixing RNA sequencing experiments with poly(dT) priming during reverse transcription (**Fig. 3B**).

Optimization of these mechanisms required a large condition screen over 186 independent experiments (**Fig. 3C; Fig. S2; Supplementary Table S1**). The following parameters and components were iteratively modified: *in vitro* RNA versus RNA within living cells, NAI-N_3_ labeling concentration and duration, bobcode sequence, bobcode nucleotide chemistry, DBCO attachment site on bobcodes, bobcode architecture, linker size and type between bobcode and DBCO, incorporation of unique molecular identifier (UMI) and primer sequences into bobcodes, and a variety of enzymatic conditions and workflow steps. We performed nanopore sequencing^36^ at shallow depth to determine barcoding accuracy of the resulting cDNA libraries, defined as the percentage of reads in which the sample-specific barcode matched the mouse or human species alignment of the insert. We set a target barcoding accuracy of >99%, more stringent than the >96% accuracy benchmark reported for DRUG-seq^10^.

These experiments resulted in a wide range of barcode accuracies, reflecting both the intrinsic limitations of each transfer mechanism and our relative success at identifying and reliably implementing protocol changes to optimize them. All transfer mechanisms yielded >90% barcode accuracy in multiple experimental conditions (**Fig. 3C**). Nevertheless, species mixing experiments in which DBCO was attached to the 3’ end of a terminal rGrGrG sequence on bobcodes (3’DBCO TSO bobcode) consistently achieved the highest barcode accuracy under the widest range of conditions and often surpassed 99.5% accuracy, the empirically determined sensitivity limit of the assay (data not shown) (**Fig. 3C**). The 3’DBCO TSO bobcode configuration also yielded >99% barcoding accuracy when reducing NAI-N_3_ RNA labeling concentrations and varying reverse transcription parameters including starting input RNA amounts and temperature between 42°C and 54°C (**Fig. S3**). Thus, we found that 3’DBCO TSO bobcodes were a reliable, robust, fast, easy to use, and versatile transfer mechanism that could achieve consistently high barcoding accuracies.

### Capturing full length RNA sequencing with bobcodes using random hexamer priming during multiplexed reverse transcription

Because bobcodes are attached along the body of RNA molecules rather than only at the 5’ or 3’ ends, we tested whether priming with random hexamers during reverse transcription would capture the full length of RNAs instead of only the 3’ ends in existing methods (**Fig. 4A**).

**Figure 4.**
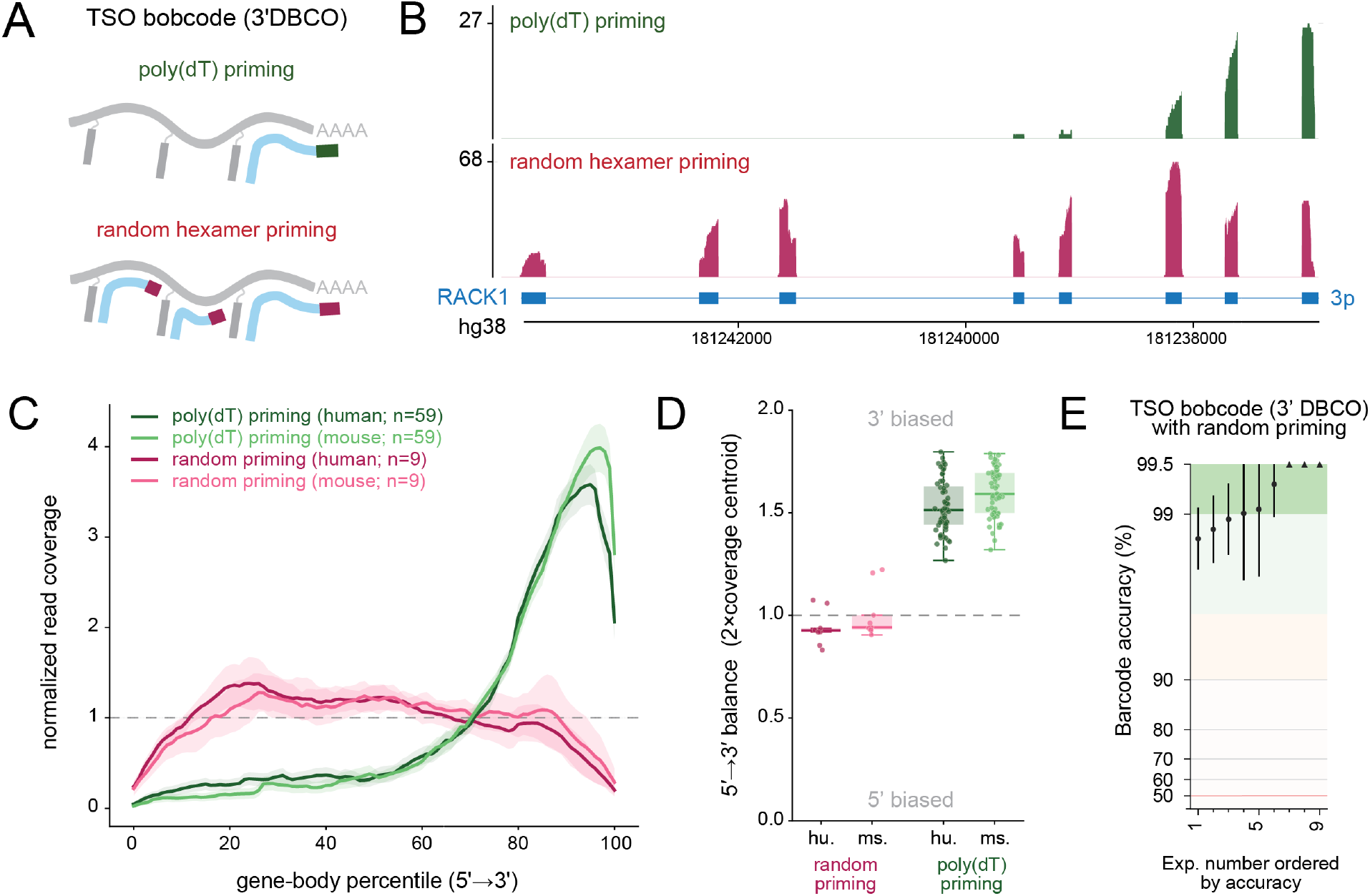
Switching from poly(dT) to random hexamer priming enables full length RNA sequencing. (A) Diagram of reverse transcription priming strategies. (B) RNA sequencing coverage demonstrating random hexamer priming captures all exons of the human RACK1 gene. (C) Transcriptome-wide gene body coverage plot profiles with PicardTools. Poly(dT) samples were taken from Fig. 3C and samples with fewer than 1000 reads were removed (n = 59 TSO bobcode (3’DBCO) experiments of 73). 95% confidence intervals are shown as light bands. (D) Calculated gene coverage balance box plot. (E) Species-specific barcoding accuracy using TSO bobcodes (3’ DBCO) after multiplexed cDNA synthesis with random priming. 95% confidence intervals shown reflecting counting error only from a single sequencing experiment each (Wilson score interval). Random hexamer priming conditions described in (**Supplementary Table S2**).

Poly(dT) priming generated 3’-biased libraries, whereas random priming produced sequencing reads covering the full length of transcripts (**Fig. 4B-D**). Library preparation workflows with random priming during reverse transcription achieved similar barcode accuracies (>99.5%) to poly(dT) priming, after optimization (**Fig. 4E; Supplementary Table S2**). The distributed position of bobcodes along the length of RNA and random priming therefore enable accurate barcoding of full-length transcripts in cDNA libraries.

### Multiplexed transcriptomic drug screen with bioorthogonal barcode sequencing (BOB-seq)

We next assessed the utility of bobcodes to detect changes in RNA isoforms in a scalable, multiplexed transcriptomic drug screen. To this end, we designed a 24-plex bioorthogonal barcode sequencing (BOB-seq v0.1) workflow in which we treated cells with small molecule and ASO drugs. We chose two small molecules with different effects on RNA isoform regulation: (1) cycloheximide (CHX), an inhibitor of translation elongation widely used to stabilize isoforms degraded by nonsense-mediated mRNA decay (NMD), which changes many isoforms at once^37^; and (2), risdiplam (Ris), an FDA-approved splicing modifier that promotes inclusion of *SMN2* exon 7 with documented off-target effects on splicing in other RNAs^7,38,39^. After drug treatment, we then bobcoded cellular RNAs with two-step chemistry and multiplexed cDNA synthesis with random priming for full length RNA capture starting with HEK293T human and RAW 264.7 mouse cells (**Fig. 5A**; **Fig. S4A-C**; **Supplementary Table S3**). Without a fragmentation or tagmentation step, cDNA libraries peaked at approximately 500bp due to bobcode spacing (**Fig. S4D**). Each read pair contains a sample-specific barcode and UMI encoded within the bobcode sequence and cDNA insert downstream of non-templated nucleotides, enabling us to use both reads from Illumina sequencing for alignment (**Fig. 5B**).

**Figure 5.**
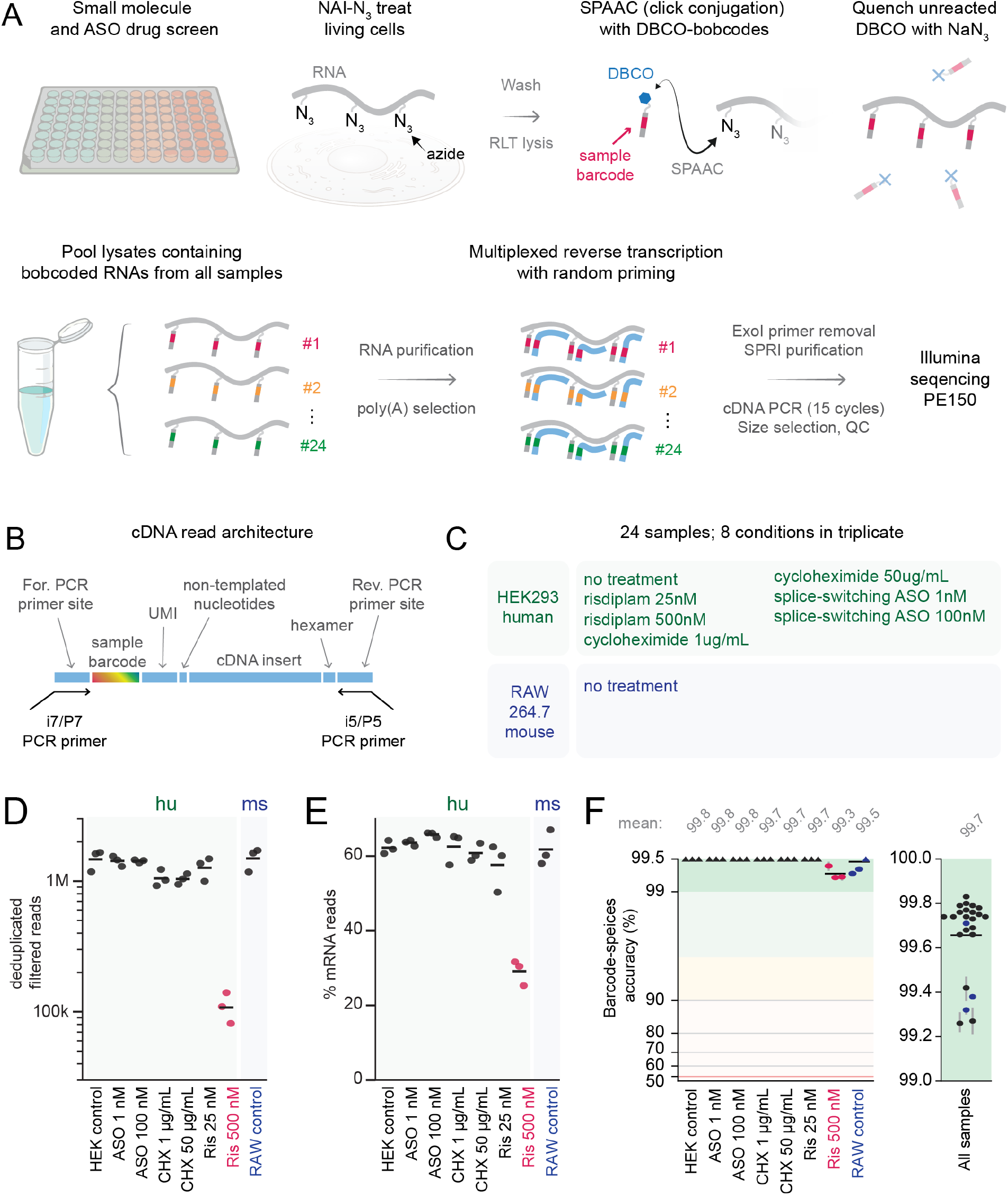
Multiplexed RNA isoform drug screen. (A) Each well of adherent cells was azide labeled, washed, lysed in RLT, bobcoded with click conjugation of DBCO to azide, and unreacted DBCO-bobcodes were quenched by spiking in sodium azide. Lysates containing bobcoded RNAs from all 24 samples were pooled prior to RNA purification, Poly(A) selection of mRNAs, and multiplexed cDNA synthesis. First strand cDNA was then prepared for Illumina sequencing. (B) cDNA read architecture and PCR priming strategy compatible with paired end 150bp (PE150) short read sequencing. (C) All sample conditions were prepared in triplicate. Three wells of RAW 264.7 mouse cells were included to assess species-specific barcode accuracy. (D) Deduplicated and quality filtered reads from each sample, presented by condition. (E) The proportion of mapped reads aligning to protein-coding mRNAs (excluding ribosomal mRNAs) from each sample. (F) Percentage of barcodes aligned to the correct species in each sample, arranged by condition.

To assess data quality, we mapped reads from all samples to the combined human and mouse transcriptomes, retained uniquely mapped reads, and collapsed PCR duplicates using each read’s UMI. TT cells treated with the highest risdiplam dose (500 nM) yielded far fewer reads, a lower fraction of mRNA reads than every other sample, and were most enriched for several genes associated with integrated stress response and cellular quiescence including PPP1R15A^40^ and DHRS2^41^, consistent with dose-dependent toxicity (**Fig. 5C–E, Fig. S4E**). We subsequently removed these samples from further analyses. Barcode to species mapping was 99.7% accurate on average across all samples, with every sample exceeding 99% (**Fig. 5F, Fig. S4F**).

We next compared BOBseq gene-expression measurements from control HEK293 cells with previously published HEK293 datasets generated by other multiplexed RNA-sequencing methods TruSeq, BRB-seq, and prime-seq^11,42–44^. Across all datasets, each sample was most similar to replicates prepared by the same method, indicating that library preparation is the dominant source of variation in these measurements (**Fig. 6A**). Enzymatic barcoding methods and BOBseq were all divergent from each other, and differential expression versus TruSeq identified that a majority of differentially expressed genes were exclusive to a specific library preparation method, with pairwise overlaps being of similar size (**Fig. 6B, Fig. S5A-C**). Together, these results indicate that random priming and chemical barcoding in BOBseq yields gene-expression measurements comparable to established RNA-sequencing methods.

**Figure 6.**
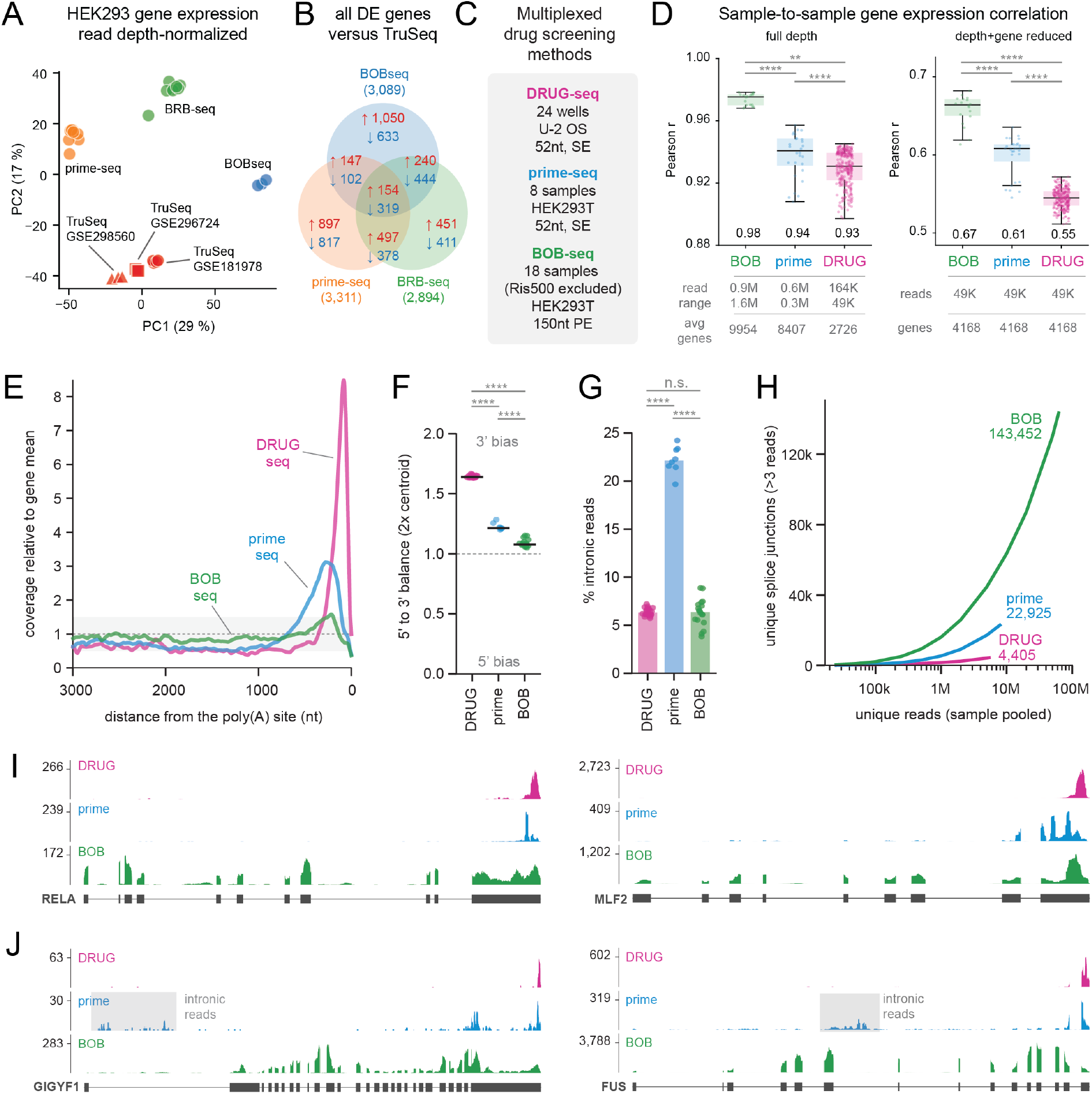
BOBseq captures full length RNA transcripts. (A) HEK293 gene expression from BOBseq (n=3 control samples) and published datasets generated from other multiplexed RNA sequencing methods. (B) Differentially expressed genes between TruSeq data and each of the other methods’ data. (C) Key information from the multiplexed drug screening method datasets used as comparators. (D) Per-replicate-pair Pearson correlation of log₂(CPM+1) across mRNA gene expression. Boxes show median and IQR; each dot is one replicate pair; the value above each box is the median r. (left) Native sequencing depth, genes with ≥5 counts in both samples. (right) All samples depth-matched with the top 4,168 co-detected genes per pair. (E) Coverage relative to the gene-body mean versus distance from the poly(A) site (transcripts ≥ 1 kb, ribosomal-protein genes excluded), showing a sharp 3′ peak for DRUG-seq and prime-seq but near-uniform coverage for BOBseq. (F) 5′-to-3′ balance (2 × coverage centroid; 1 = even) per sample. (G) Intronic reads as a percentage of mapped reads per sample. (H) Unique splice junctions (> 3 deduplicated reads) versus pooled depth. (I) Pooled, depth-matched coverage tracks for RELA and MLF2 (hg38). Y-axis reflects read coverage. (J) Same tracks for GIGYF1 and FUS, with grey shading marking prime-seq intronic reads (hg38). Statistics: (D) Brackets: two-sided Mann–Whitney U on per-sample mean correlation (the ∼independent unit; ** p<0.01, **** p<1×10⁻⁴) (F,G): Kruskal–Wallis with BH-corrected pairwise post-hoc; **** P < 10⁻⁴, n.s. = not significant. DRUG (n=24), prime (n=8), BOB (n=18; Ris500 excluded).

### Higher replicate correlation in BOB-seq gene expression measurements than existing multiplexed transcriptomics methods

Sample-to-sample variance in gene-expression measurements can be strongly influenced by technical variability in library-preparation workflows that process samples in parallel rather than as a pool. Since RNA samples in BOBseq share the same vessel for all enzymatic processing steps, we hypothesized BOBseq would show reduced gene expression variance compared to existing multiplexed transcriptomic well-based drug screening methods prime-seq (performed on HEK293 cells) and DRUG-seq (performed on U2OS cells) in which reverse transcription-based barcoding is performed individually on each sample before pooling^8,11^ (**Fig. 6C**). We found that BOB-seq exhibits significantly higher gene expression correlation and lower variance (root mean squared error (RMSE)) among replicate pairs compared to the other multiplexed well-based preparation methods (**Fig. 6D, Fig. S6A**). We observed the same result when using complete datasets generated with each method and after down-sampling read counts from BOB-seq and prime-seq to match the lower sequencing depth and gene detection of DRUG-seq samples (**Fig. 6D**). Thus, early multiplexing of samples in BOBseq prior to reverse transcription is associated with reduced sample-to-sample variance in RNA measurements.

### BOB-seq outperforms DRUG-seq and prime-seq on RNA full length and splice junction sequencing

We compared BOB-seq alignment metrics from 18 HEK293T samples (excluding the 3 high dose risdiplam samples) to DRUG-seq and prime-seq^10,11^ (**Fig. S6B-E**). Read coverage across gene bodies was most severely 3’ biased in DRUG-seq and was most even in BOB-seq (**Fig. 6E-F, Fig. S6F**). Compared to both BOB-seq and DRUG-seq, prime-seq captured nearly a 4 times higher proportion of intronic reads (**Fig. 6G**), which can complicate measures of gene expression^45^. BOB-seq captured a higher percentage of reads mapping to protein coding mRNAs than both other methods (**Fig. S6B**). Reflecting more complete end-to-end RNA transcript capture, BOB-seq captured nearly 25 times more splice junctions than DRUG-seq and 6 times more than prime-seq (**Fig. 6H**). We examined read coverage across individual genes and confirmed BOB-seq often captures every exon and splice junction in highly expressed genes (**Fig. 6I-J**). In contrast, reads from DRUG-seq and prime-seq were almost exclusively aligned to the 3’ end of genes with high coverage in intronic regions specifically in prime-seq (**Fig. 6J**). Thus, BOB-seq’s use of chemically conjugated barcodes along RNAs and random priming enables the capture of RNA isoforms and RNA splicing events that are missed by enzymatic 3’ end-only barcoding methods.

### Identification of gene expression and RNA splicing changes with drug treatment

We examined gene expression in our BOB-seq samples by PCA and determined CHX drove the largest changes in gene expression in a dose dependent manner (**Fig. 7A**). CHX dose-dependently increased the expression of JUN, consistent with prior reports of CHX induction (**Fig. 7B, Fig. S7A**)^37^. We next examined CHX dose-dependent effects on RNA isoforms by examining differentially spliced junctions compared to the control condition (**Fig. 7C**). Among the top affected genes was HRAS which has a known poison exon containing a premature termination codon that triggers NMD^46^. Next, we examined the effects of 25nM risdiplam on gene expression and RNA isoforms and found minimal changes passing significance across the transcriptome, consistent with low off target effects at this dose (**Supplementary Table S4-7**). We performed a custom read alignment to exon 7 of the SMN1 and SMN2 genes to accurately identify reads carrying the one nucleotide difference that distinguishes the two genes. In 25nM risdiplam treated samples, the proportion of reads skipping exon 7 were reduced, consistent with risdiplam’s known RNA splicing regulatory effects on SMN2 splicing for the treatment of SMA (**Fig. 7D, Fig. S7B**). We examined gene coverage on differentially spliced genes AKAP8 and CLASRP in CHX treated samples and confirmed the change in splice site selection in low and high dose conditions (**Fig. 7E**). We also examined gene coverage of FOXM1 and MADD, previously identified off targets of risdiplam^38,47,48^, and identified changes in RNA splicing in the 25nM risdiplam condition compared to controls (**Fig. 7F**).

**Figure 7.**
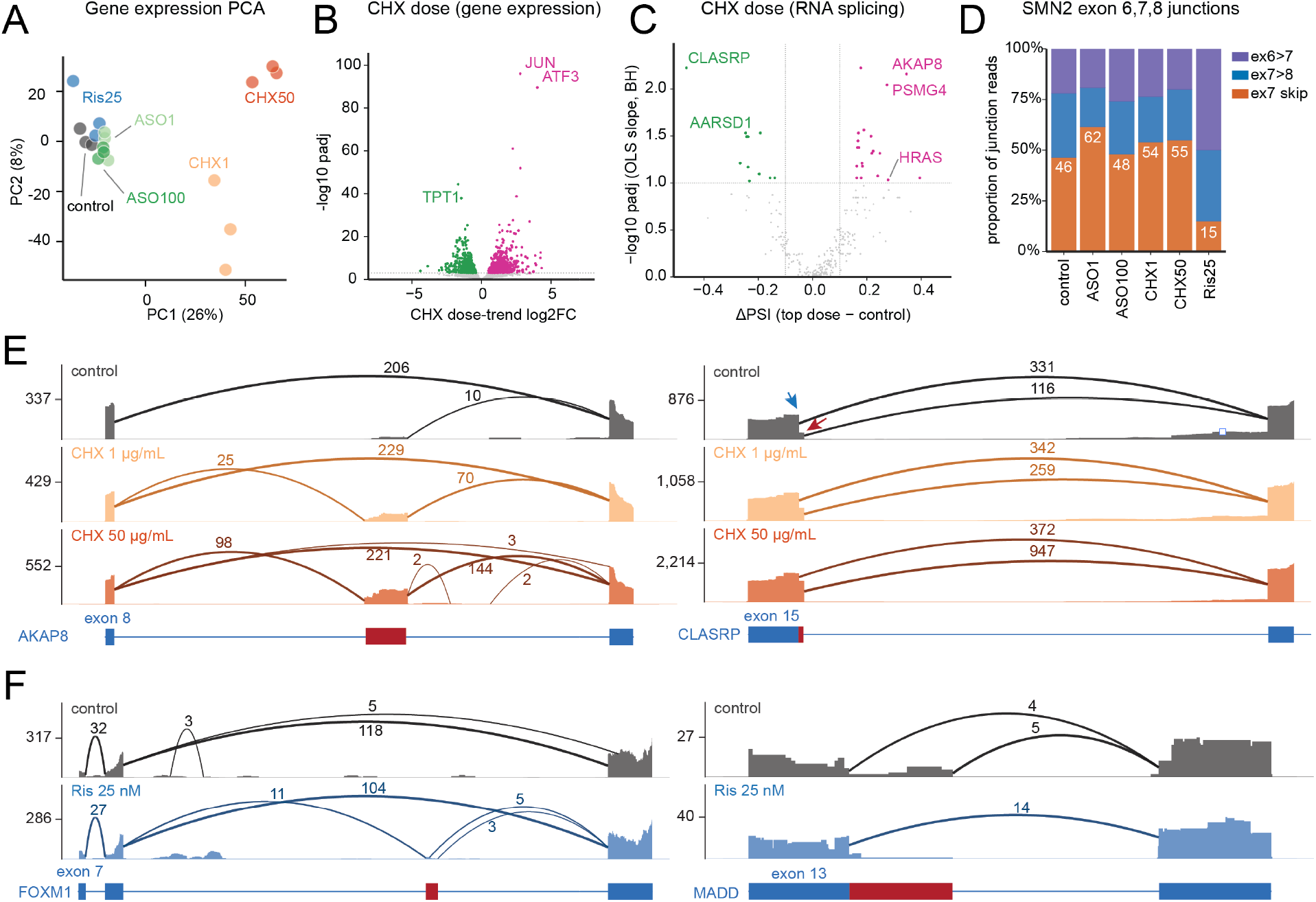
BOB-seq multiplexed transcriptomic drug screen. (A) Principal components analysis of gene expression from BOBseq samples, excluding Ris500. (B) CHX dose-dependent changes in gene expression. (C) CHX dose-dependent changes in percent spliced in (PSI) values for splice junctions. (D) Quantification of junction reads for SMN2 exon 7. (E, F) Gene coverage plots (hg38) for selected genes identified as exhibiting differential RNA splicing after CHX treatment (E) and previously identified splicing off-targets of risdiplam (F). Y-axis reflects read coverage, and numbers over arcs reflect splice junction reads. Red bars on gene body plots reflects alternative exon boundaries.

### BOB-seq workflow is shorter and less complex compared to existing multiplexed bulk RNA sequencing methods

We evaluated BOB-seq workflow characteristics compared to existing methods based upon protocol descriptions^9–11^. Of note, the unique reagents in BOB-seq include DBCO conjugated to oligonucleotides and the chemical NAI-N_3_, both of which are affordable and easily available. No other specialized reagents or equipment is required to perform BOB-seq.

Protocol duration in DRUG-seq and prime-seq is affected by the need for cDNA tagmentation and enzymatic fragmentation, respectively (**Fig. S7C**). However, in BOB-seq, bobcode spacing sets cDNA size directly without the need for fragmentation or tagmentation. Ideal for paired end 150bp (PE150) short read sequencing, BOB-seq libraries peaked at ∼500 bp (∼350 bp insert and ∼150bp adapters) after cDNA amplification but before double sided-SPRI size selection (**Fig. S4D**).

Though we had designed the BOB-seq v0.1 protocol for data quality and not for protocol speed, we estimated that BOB-seq requires approximately 95 minutes less incubation time than DRUG-seq and prime-seq^10,11^. BOB-seq v0.1 has fewer steps and involves the least per-sample handling times prior to sample pooling (**Fig. S7C-D**). Prime-seq, the next shortest method, requires processing each sample with complex workflow steps including per-well proteinase K, bead cleanup, and DNase I. Thus, the improvements in data quality and full-length RNA capture with BOB-seq are associated with less protocol time and complexity, rather than more.

PCR chimeras between barcoded molecules can drive barcode hopping and barcode accuracy is better preserved if amplification cycle number is kept low^11,49,50^. While BOB-seq amplifies cDNA in a single 15-cycle PCR, a total of up to 33 cycles in DRUG-seq and 28 cycles in prime-seq across two pre-fragmentation and post-fragmentation PCR amplification steps are reported in these methods. BOB-seq’s 0.3% barcode swapping rate (100 - 99.7% barcode accuracy) is more than 10-fold lower than the 4% rate (100 – 96% barcode accuracy benchmark) reported in DRUG-seq and lower than the ∼0.5% barcode swapping rate reported in prime-seq^10,11^. Because BOB-seq has 18 fewer PCR cycles than DRUG-seq (up to ∼262,144-fold less amplification), we hypothesize it is less likely to exhibit amplification biases in sequenced data^51,52^.

## Discussion

Multiplexed sequencing technologies are the basis of scalable transcriptomic measurement^53,54^. Barcoding and pooling methods exponentially cut per-sample cost and labor, enabling the creation of single-cell atlases across human organs, genetic perturbation experiments probing gene regulatory pathways, and compound screens for drug discovery^8,10,55–60^. A single researcher can now profile more than thousands of cells in a single experiment with only modestly more wet lab effort than it takes to prepare a single sample^59,61^.

Yet scale has come at a severe cost to the richness and reliability of each sample’s measurement^62–65^. This is particularly true for the capture of RNA isoforms, which drive diversification of the proteome, alter gene-regulatory networks, fuel evolutionary innovation, and drive the pathophysiology of many diseases^66–69^. Because every existing multiplexed RNA sequencing method only attaches a single barcode to the 5′ or 3′ end of a molecule, isoform information far from the barcode is lost when libraries are fragmented for short-read sequencing^8–11,55,70–72^.

Furthermore, barcoding is often inaccurate in many workflows due to barcode contamination, especially in single cell methods that contain large fractions of ambient RNAs released from lysed cells^10,73,74^. Consequences can be severe, including mistaken attribution of cell identities^75^. Thus, both the resolution and accuracy of RNA measurements are often compromised at scale.

To restore data quality in scalable transcriptomics, we show that bioorthogonal barcodes (“bobcodes”) can be covalently conjugated to native RNA at multiple internal positions by acylation and click chemistries. Compared to existing well-based drug screening methods, bobcodes improve barcoding accuracy, capture full-length RNA isoform information, and reduce workflow complexity by eliminating tagmentation and fragmentation steps and pre-amplification PCR. Because bobcoded RNAs are pooled earlier than in any existing workflow method, sample-to-sample variability and workflow complexity caused by parallel reactions is reduced.

To the best of our knowledge, our method is the first to enable multiplexing of RNA sample processing before reverse transcription or any other enzymatic step (**Fig. 8**). With poly(A) mRNA selection of pooled bobcoded RNAs and random-primed multiplexed cDNA synthesis, we demonstrate that bobcodes enable a transcriptome-wide screen of drug effects on RNA isoforms. Though we performed our screen on 24 samples, it can be easily scaled to larger number of samples by expanding the number of unique barcode sequences.

**Figure 8.**
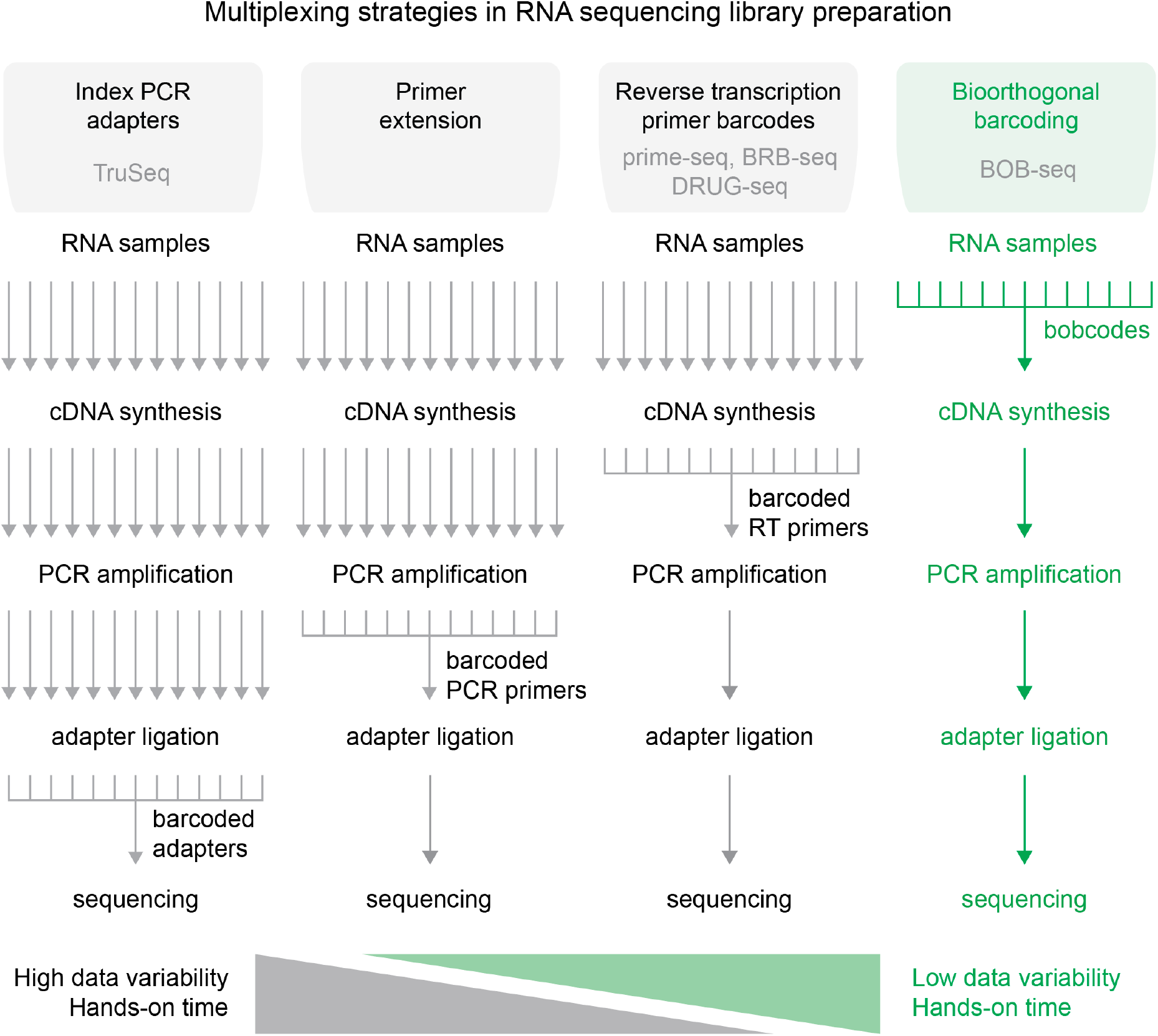
Diagram of multiplexing strategies in nucleic acid sequencing workflows. Bobcoding uniquely enables barcoding and pooling of RNA samples prior to any enzymatic step, reducing hands-on-time and sample-to-sample variability caused by parallel processing.

Beyond the well-based screening assay described herein, chemoselective conjugation chemistry applied to nucleic acid barcoding represents a new platform for multiplexed sequencing. In principle, the same two-step chemical barcoding method is adaptable to single cell combinatorial indexing, droplet-based barcode installation, and spatially resolved barcoding in tissue sections^76,77^. Because of the fidelity and robustness of the chemistry used and ability to place multiple barcodes on an individual mRNA, we predict that RNA capture rates could be significantly improved in single cell measurements using bobcodes. By collapsing per-sample processing into a single pooled workflow, bobcodes offer a potentially lower-cost, lower-labor, and lower-variability alternative to gene expression profiling methods that still prepare samples in parallel enzymatic reactions, such as qPCR.

Lastly, existing transcriptomic methods struggle to capture RNA sequencing data from degraded or FFPE material because in situ reverse transcription barcoding with poly(dT) primers is unreliable on crosslinked nucleic acids^78–80^. Bobcoding has the potential to avoid these limitations, and we anticipate its compatibility with multiplexed de-crosslinking and rRNA depletion methods for degraded samples prior to cDNA synthesis. We believe chemical barcoding has the potential to offer greater versatility in experimental conditions for transcriptomics.

By bringing isoform-level resolution to highly multiplexed RNA sequencing without sacrificing scale, bobcodes can make large transcriptomics screens more informative for splicing biology, accelerate drug discovery, and enable the generation of more accurate training data for machine-learning models.

## Limitations

As presented, BOB-seq has the following practical and technical limits in the version 0.1 described. First, removing excess NAI-N_3_ requires washing cells after labeling, which may cause cell loss when working with suspension cells or poorly adherent cells and may limit applicability to some cultured cell types. Second, the fraction of input mRNA molecules converted to cDNA should be benchmarked against existing methods with reference standards, for example with ERCC spike-ins. Third, comparison metrics of BOB-seq with other transcriptomic methods were performed on previously published datasets and our present analyses are limited by low number of replicates. We were also not able to independently verify published methods’ barcoding accuracy since experimental data required for analysis was not deposited. Finally, because isoforms are reconstructed from short inserts rather than long inserts end-to-end as in long-read sequencing, distant splicing events on the same molecule cannot be directly phased.

## Author contributions

N.D.A. invented the technologies described, performed pilot studies, developed analysis pipelines, and led the investigation. T.N.M., J.B., and A.K. contributed to study design and performed experiments and optimizations. N.D.A., B.N., and M.A. performed computational analyses. N.D.A. supervised the study and acquired funding. N.D.A. wrote the manuscript and all the authors reviewed and approved the manuscript.

## Competing interests

Sculpta, Inc. has filed provisional patent applications covering chemical barcoding, multiplexed cDNA synthesis, and their applications in nucleic acid measurements. All authors are employees, contractors, and/or shareholders of Sculpta, Inc. N.D.A. is the founder and CEO of Sculpta, Inc.

## Funding

This work was privately funded. No government, university, or non-profit funding directly contributed to this work.

## Artificial Intelligence Use Statement

We used large language models from Anthropic (Opus, Fable) and OpenAI (GPT4, GPT5) for coding, research, bioinformatics, and experimental design. The main text and legends were written by humans, though language models were used for revising and editing. The figures were made by humans. The scientific inventions and technologies described herein were invented by humans alone, not by artificial intelligence.

## Data and Code Availability

Sequencing data have been deposited in the NCBI SRA under BioProject PRJNA1532586 (SRA study SRP738601) which will be released September 2026. Code is available at https://github.com/Sculpta/bobseq (release v0.1, MIT license). The Patchwork (Sculpta, Inc.) RNA splicing bioinformatics software is proprietary and is not publicly available.

## Key Reagent List

| Reagent | Vendor | Catalog Number |
| --- | --- | --- |
| NAI-N3 (methyl 2-(azidomethyl)imidazo[1,2-a]pyridine-6-carboxylate) | Enamine | EN300-28312133 |
| DBCO-PEG4-NHS ester, (Dibenzocyclooctyne-PEG4-N-hydroxysuccinimidyl ester) | Millipore Sigma | 764019 |
| 7.5% Sodium bicarbonate | Millipore Sigma | S8761 |
| DMSO (Dimethyl sulfoxide) | Millipore Sigma | D2438 |
| Azide agarose | Millipore Sigma | 900957 |
| Tween-20 | Millipore Sigma | P1379 |
| Sodium chloride solution | Millipore Sigma | S5150 |
| UltraPure™ 0.5M EDTA, pH 8.0 | ThermoFisher | 15575020 |
| HEPES, 1.0M buffer soln., pH 7.0 | ThermoFisher | J62688.AK |
| DPBS, calcium, magnesium | ThermoFisher | 14040133 |
| Bovine Serum Albumin | R&D Systems | 5217 |
| Oligo Clean & Concentrator | Zymo | D4060 |
| RNeasy Mini Kit | Qiagen | 74104 |
| Buffer RLT | Qiagen | 79216 |
| HEK293T cell line (human) | ATCC | CRL-3216 |
| RAW 264.7 cell line (mouse) | ATCC | TIB-71 |
| Dulbecco's Modified Eagle Medium (DMEM, 1X) | ThermoFisher | 11965092 |
| HyClone USDA import tested Fetal Bovine Serum (FBS) | VWR | 89133-094 |
| Penicillin-Streptomycin (5,000 U/mL) | ThermoFisher | 15070063 |
| TrypLE Express Enzyme (1X), no phenol red | ThermoFisher | 12604013 |
| Poly-D-Lysine | ThermoFisher | A3890401 |
| Risdiplam | MedChemExpress | HY-109101 |
| Cycloheximide, ultrapure, 95% | ThermoFisher | J66901.03 |
| Opti-MEM Reduced Serum Medium | ThermoFisher | 31985070 |
| Lipofectamine 3000 Transfection Reagent | ThermoFisher | L3000008 |
| Ethanol, 200-proof, molecular biology grade | Sigma | E7023-1L |
| CleanNGS DNA & RNA Clean-Up Magnetic Beads | Bulldog Bio | CNGS050 |
| Sodium azide, 99% | ThermoFisher | 14314.09 |
| Nuclease-free water (not DEPC-treated) | ThermoFisher | AM9939 |
| NEBNext Poly(A) mRNA Magnetic Isolation Module | NEB | E7490S |
| Maxima H Minus Reverse Transcriptase (200 U/μL) | ThermoFisher | EP0751 |
| Deoxynucleotide (dNTP) Solution Mix | NEB | N0447S |
| Exonuclease I (E. coli) | NEB | M0293S |
| iTaq Universal SYBR Green Supermix | Bio-rad | 1725124 |
| KAPA HiFi HotStart plus dNTPs, 250U | Roche | 7958897001 |
| High Sensitivity RNA ScreenTape | Agilent | 5067-5579 |
| High Sensitivity RNA ScreenTape Sample Buffer | Agilent | 5067-5580 |
| High Sensitivity RNA ScreenTape Ladder | Agilent | 5067-5581 |
| Optical Tube Strips (8x Strip) | Agilent | 401428 |
| Optical Tube Strip Caps (8x Strip) | Agilent | 401425 |
| High Sensitivity DNA ScreenTape | Agilent | 5067-5584 |
| High Sensitivity D1000 reagents | Agilent | 5067-5585 |
| E-Gel Agarose Gels with SYBR Safe DNA Gel Stain | ThermoFisher | A42135 |
| 100bp DNA Ladder | NEB | N3231L |

## Methods

### Dose-dependent NAI-N₃ labeling of RNA

Total RNA was extracted from THP-1 cells (ATCC, TIB-202) with the RNeasy Mini Kit (Qiagen, 74104). Acylation reactions (10 µL) contained 100 ng total RNA in 50 mM HEPES pH 7.4 and NAI-N₃ (MedChemExpress, HY-103006; 1 M in anhydrous DMSO) at 0, 1.25, 5, 20, 80, 100 or 125 mM, and were incubated at 37 °C for 10 min. Azide-labeled RNA was purified (Monarch RNA Cleanup Kit, New England Biolabs, T1030) and eluted in 10 µL water. For strain-promoted azide–alkyne cycloaddition, 8 µL of azide-labeled RNA was reacted with 0.1 mM DBCO-FAM (Broad Pharm, BP-28970) in 10 µL 50 mM HEPES pH 7 at 37 °C for 30 min; unreacted probe was removed by RNA cleanup (New England Biolabs, T1030) and RNA was quantified by A₂₆₀ (NanoDrop One, Thermo Fisher). Conjugate fluorescence was measured on a Varioskan LUX reader (Thermo Fisher; excitation 490 nm, emission 520 nm) against a DBCO-FAM standard curve in PBS pH 7.4 (Gibco, 10010049). Label density was expressed as FAM molecules per 1,000 nucleotides, calculated from the standard-curve-derived FAM amount and the nucleotide amount inferred from RNA mass.

### Effect of NAI-N₃ labeling on cDNA library insert size

Total RNA was labeled with NAI-N₃ (100 ng per reaction) and purified. Where indicated, azide-labeled RNA was conjugated to 0.1 mM DBCO-FAM (Broad Pharm, BP-28970) in 50 mM HEPES pH 7.4 at 37 °C for 30 min and purified twice (Monarch RNA Cleanup Kit, New England Biolabs, T1030). Reverse transcription was performed with SuperScript IV (Thermo Fisher, 18091050) in 5 µL reactions containing ∼10 ng RNA, 10 µM poly(dT)-VN RT primer (5′-ATACTCGTGACATCAGCTTCCT₃₀VN-3′; IDT), 1 µM biotinylated template-switching oligonucleotide (5′-biotin-GCTAATCATTGCAAGCAGTGGTATCAACGCAGAGTACATrGrGrG-3′; IDT), dNTPs (Sigma-Aldrich, 71004M), SuperScript IV buffer and DTT, with incubation at 23 °C for 10 min then 50 °C for 10 min. cDNA was purified (Monarch PCR & DNA Cleanup Kit, New England Biolabs, T1030) and amplified with Q5 High-Fidelity 2× Master Mix (New England Biolabs, M0492) using primers against the RT-primer (5′-ATACTCGTGACATCAGCTTCC-3′) and TSO (5′-GCAGTGGTATCAACGCAG-3′) handles (98 °C, 30 s; then 20 cycles of 98 °C 10 s, 65 °C 20 s, 72 °C 2 min). Libraries were purified (Monarch PCR & DNA Cleanup Kit, New England Biolabs, T1030) and sized on a 4200 TapeStation with D5000 ScreenTape (Agilent, 5067-5588).

### Dose-dependent RNA barcoding quantified by gel shift

Total RNA (isolated as above) was fragmented in 100 mM Tris-HCl pH 7.0, 150 mM KCl, 6 mM MgCl₂ at 94 °C for 5 min and purified (Monarch RNA Cleanup Kit, New England Biolabs, T1030). Acylation reactions (10 µL) contained 100 ng fragmented RNA and NAI-N₃ (MedChemExpress, HY-103006) at 0, 20, 80 or 200 mM in 50 mM HEPES pH 7.4, and were incubated at 37 °C for 10 min; azide-labeled RNA was purified as above and eluted in 10 µL water.

A DBCO-functionalized barcode oligonucleotide was prepared from a 51-nt oligonucleotide bearing a 5′ amino-C6 linker (5′-ACACTCTTTCCCTACACGACGCTCTTCCGATCTTCAGCAGTACCTGGTAGC-3′; IDT). The 5′ amine was conjugated to DBCO-PEG4-NHS ester by combining 200 µL of 100 µM oligonucleotide (nuclease-free water), 54 µL 7.5% sodium bicarbonate pH 8.3, 10 µL of 25 mM DBCO-PEG4-NHS ester (MedChemExpress, HY-140272) in DMSO and 20 µL DMSO (284 µL total), for 1 h at room temperature protected from light. The DBCO-barcode oligonucleotide was purified (Monarch Oligonucleotide Cleanup Columns, New England Biolabs, T1040), eluted in 100 µL nuclease-free water, quantified by A₂₆₀ (NanoDrop One, Thermo Fisher) and stored at −20 °C.

For conjugation, 8 µL of azide-labeled RNA was reacted with 0.1 mM DBCO-barcode oligonucleotide in 1.0X SPRI at 37 °C for 30 min. Unconjugated oligonucleotide was removed by SPRI and RNA cleanup (New England Biolabs, T1030) and 10 ng of barcoded RNA was analyzed on a 4200 TapeStation with RNA ScreenTape (Agilent, 5067-5576). Barcode-coupling density was inferred from the apparent size shift relative to unlabeled RNA.

### Bobcode transfer mechanism screen

Over 186 experiments, a variety or parameters and components were changed to engineer a library workflow method that consistently achieved greater than 99% barcoding accuracy in species mixing experiments. Cell lines used in these experiments included THP-1 cells (ATCC, TIB-202), HEK293T cells (ATCC #CRL-3216), and RAW 264.7 cells, and in some experiments, RNA was isolated from cells and purified prior to azide labeling. We used Maxima H Minus Reverse Transcriptase for these studies. The specific conditions and variants are described in Supplemental Fig X. We used nanopore sequencing to rapidly capture shallow sequencing depths to assess barcode accuracy and other parameters.

cDNA libraries were multiplexed with the Native Barcoding Kit 24 V14 (SQK-NBD114.24; Oxford Nanopore Technologies) and sequenced on a MinION Mk1D with R10.4.1 flow cells. Reads were basecalled and demultiplexed in MinKNOW with the high-accuracy model dna_r10.4.1_e8.2_400bps_hac@v5.2.0. Each native barcode corresponded to one library, and all passing reads of each library were analysed, with three exceptions. For three libraries sequenced in a 22-hour run (0.94–1.43 million reads each), only the first 10,000 reads of each library were analyzed. Where the same libraries had been sequenced in two runs, the reads of both runs were pooled (15 libraries from two sets). For one set of five libraries sequenced in three runs, a single run was analyzed. Read identifiers were unique within every library.

Each library contained human and mouse material carrying two different bobcodes, one assigned to each species. The bobcode lies immediately 3′ of a constant 12-nt sequence (the anchor) within an oligonucleotide whose structure depends on the chemistry. In poly(dT) priming, a 6-nt code (12 nt in one 90-mer design) is part of the oligo(dT) reverse-transcription (RT) primer. In splint-mediated ligation, a 4-nt code (8 nt in the BOB-A design) is carried by a primer ligated to the RT primer through a splint oligonucleotide. In the TSO bobcode (5′DBCO) chemistry, a 7-nt code follows a Nextera Read 1 backbone in the template-switching oligonucleotide (TSO), or a 4-nt code follows a SMARTer backbone in one design. In the TSO bobcode (3′DBCO) chemistry, a 7-nt code follows a SMARTer TSO backbone or a TruSeq Read 1 handle. The construct and the two codes of every library are listed in Supplementary Table S1; the anchor of each construct is given in the code repository. Read processing and alignment

Native barcode and adapter sequences were removed from both ends of each read. At the 5′ end, everything up to and including the barcode flank CAGCACCT, searched within the first 80 nt, was removed; at the 3′ end, everything from the last occurrence of its reverse complement AGGTGCTG within the final 80 nt onwards was removed. Trimmed reads were aligned with STARlong 2.7.11b ^81^ to a combined human and mouse genome built from the Ensembl ^82^ GRCh38 and GRCm39 primary assemblies, with chromosome names prefixed by species, and Ensembl release 113 gene annotation for both species. The index was built with a splice-junction database overhang of 99 nt and a dense suffix array. Each library was aligned separately with identical settings, and all alignments of each read were reported. The alignment parameters were:

--outFilterMultimapNmax 10000

--outFilterScoreMinOverLread 0.3

--outFilterMatchNminOverLread 0.3

--outFilterMismatchNmax 1000

--outFilterMismatchNoverLmax 0.3

--alignIntronMax 1000000

--alignSJDBoverhangMin 1

--alignEndsType Local

--seedPerReadNmax 100000

### Read classification

A read was assigned to human or mouse only if all of its alignments (primary, secondary and supplementary) lay on that species’ chromosomes. Reads with alignments on both genomes were classed as ambiguous and were not scored.

Each mapped read was assigned to one RNA category by the first matching rule: (i) rRNA, if any alignment overlapped one of 99 rDNA loci (provided with the code); (ii) mitochondrial, if the primary alignment was on the mitochondrial chromosome; (iii) rRNA, if the primary alignment overlapped an exon of a gene with an rRNA, Mt_rRNA or rRNA_pseudogene biotype; (iv) other, if the primary alignment overlapped an exon of a ribosomal-protein gene (protein-coding genes named RPL, RPS, MRPL or MRPS, excluding the RPS6K kinases; 342 genes across the two species); (v) mRNA, if the primary alignment overlapped an exon of any other protein-coding gene; and (vi) other, for all remaining reads, including lncRNA and other non-coding, intronic and intergenic reads. Exon overlap was assessed on the aligned blocks of the primary alignment, excluding splice gaps and regardless of strand.

Reads whose aligned bases had a normalized dinucleotide Shannon entropy below 0.65 (entropy divided by its 4-bit maximum; soft-clipped bases excluded) were classed as low-complexity and were not scored. This removes reads whose species assignment rests on homopolymers or simple repeats.

### Barcode calling

The species code was read from the bases immediately 3′ of an exact match to the library’s 12-nt anchor, searched on both strands. A barcode unit was defined as an anchor followed by a sequence within one mismatch of either of the library’s two codes; anchor matches less than 27 nt apart on the same strand were counted as one unit. Reads with no unit, or with two or more units (chimeras, including reads carrying both codes), were not scored. In reads with a single unit, the code had to match one of the two codes exactly, with no mismatches or indels. The one-mismatch tolerance was used only to detect units and never made a read scorable. Codes belonging to other libraries were ignored.

### Barcode accuracy and statistics

A read was scored if it was assigned to a single species, was classified as mRNA, was not low-complexity and carried a single exact code. A scored read was correct if the species assigned to its code matched the species of its alignment. Barcode accuracy was calculated as the number of correct reads divided by the number of scored reads, with a 95% Wilson score interval ^83^. Accuracy was also calculated separately for reads aligned to human and to mouse. The intervals reflect counting error only and do not include variation between libraries or experiments.

For each library, the table of codes against aligned species was inspected. Where the reads carrying a code (codes with at least 20 scored reads) did not align mainly to that code’s declared species, this was noted in Supplementary Table S1; accuracy was reported for every library. Libraries with fewer than 100 scored reads were excluded (11 libraries), as were designed negative controls, single-species controls and single-code designs, for which accuracy cannot be defined. Excluded libraries and the reasons for exclusion are listed in Supplementary Table S1.

The analysis comprised 186 libraries: 59 poly(dT) priming, 34 splint-mediated ligation, 20 TSO bobcode (5′DBCO) and 73 TSO bobcode (3′DBCO). In total, 512,565 reads were scored (median 1,639 per library).

### Code collisions

In two constructs, one species code is identical to the sequence at the code position of the standard, unbarcoded SMARTer TSO (…CAACGCAGAGTACATrGrGrG): the human code AGTACAT of the SMARTer-backbone 7-nt TSO (3′DBCO) and the mouse code TACA of the 4-nt design (5′DBCO). Products of an unbarcoded TSO would therefore be counted as carrying that code. Accuracy of the SMARTer-backbone construct (median 99.4%, 28 libraries) was not lower than that of the TruSeq Read 1 construct, which has no such collision (median 99.0%, 45 libraries). The 4-nt design contributed one library.

### Software

The analysis used Python 3.12 (standard library only), STARlong 2.7.11b and samtools 1.20 ^84^. The figure and supplementary table were produced with matplotlib 3.10 and openpyxl 3.1.

### Data and code availability

Raw nanopore reads for the 186 libraries (one FASTQ file per library, with a manifest of read counts, checksums and source files) will be deposited in a public repository. The accompanying documentation describes how the MinKNOW output files were joined, which libraries were pooled or subsampled, and which sequenced libraries were not analyzed. The complete analysis code, including the processing program, the general rules and per-run table, the rDNA loci, the reference-building instructions with contig checksums, and the figure and supplementary-table scripts, is released under the MIT license by Sculpta, Inc. It reproduces the figure and Supplementary Table S1 from the deposited reads.

### Random priming

#### Experimental iterations

As with the polydT experimental iteration screen, we changed conditions and starting RNA materials to be used with Maxima H Minus Reverse Transcriptase and either 6N or 9N random primers with a 5’ primer handle for subsequent PCR amplification.

#### Sequencing and alignment

Random-priming 3′TSO bobcode libraries (bob-v1 construct; random N-mer RT primer in place of oligo-d(T)) from human/mouse species-mixing experiments were sequenced on Oxford Nanopore. Reads were trimmed of the ONT/native-barcode flank and aligned with STARlong (v2.7.11b) to a combined *Homo sapiens* GRCh38 + *Mus musculus* GRCm39 genome (Ensembl release 113) whose contigs were prefixed HUMAN_/MOUSE_, so that read species was assigned by chromosome of alignment; BAMs were coordinate-sorted and indexed with samtools (v1.20). A single dense index was used and one STARlong process was run at a time.

#### Barcode accuracy

For each barcode, species-unique protein-coding (mRNA) reads passing low-complexity, single-barcode-unit and exact-code filters were scored, and a read was called correct when the species encoded by its 7-mer bobcode matched the species of its alignment; ribosomal-RNA and mitochondrial reads were excluded. Accuracy was reported on all scored mRNA reads with a 95% Wilson confidence interval (reads capped at 10,000 per sample) and plotted per experiment ranked by accuracy.

*Gene-body (5′→3′) coverage — Picard profile*.

Per-transcript-normalized coverage was computed with Picard CollectRnaSeqMetrics (v3.4.0, STRAND_SPECIFICITY=NONE), run once against a HUMAN_- and once against a MOUSE_- prefixed refFlat so that human- and mouse-gene metagenes were obtained from the same alignment. Gene models were protein-coding Ensembl-113 transcripts with rRNA, mitochondrial and ribosomal-protein genes excluded. Before quantification, alignments were restricted to primary records with aligned cDNA length ≤500 bp (Σ CIGAR M/I/=/X), and MINIMUM_LENGTH was set to 1000 so only transcripts ≥1000 bp contributed. Profiles are the position-wise mean of per-sample normalized-coverage histograms (101 bins) with a 95% confidence interval (t-distribution); a sample was included only if both its human and mouse tracks had ≥1000 scored reads.

*5′→3′ balance (bar plot)*.

For each sample and species, a scalar balance was defined as twice the coverage centroid of the gene-body metagene (2·Σpᵢcᵢ/Σcᵢ, pᵢ∈[0,1]): 1 = evenly balanced, <1 = 5′-leaning, >1 = 3′-leaning. Per-sample values were shown as box plots (poly(dT) vs random priming, human and mouse).

#### Single-gene coverage (RACK1 genome-browser plot)

Per-base coverage over RACK1 was computed from primary alignments with aligned cDNA length ≤500 bp and displayed for two representative samples — a poly(dT) library (run 260730_RT_conditions_BC11-13, barcode12) and a random-priming library (run 260827_BC14_16, barcode15) — against all annotated RACK1 transcript isoforms (Ensembl 113), coloured by transcript biotype.

### 24-plex RNA isoform screen

#### Bobcode preparation by DBCO-PEG4-NHS coupling

133 μM of custom synthesized BOBcode (IDT) was mixed with 180 mM sodium bicarbonate (Qiagen, S8761), 2.6% DMSO (Millipore Sigma, D2438), and 2.6 mM DBCO-PEG4-NHS (Millipore Sigma, 764019) ester. This was incubated at room temperature for 1 hour, protected from light. The solution was then purified using the Zymo Oligo Clean & Concentrator Kit (Zymo D4060) following manufacturer protocols and eluted in in nuclease free (ThermoFisher, AM9939) water to get a final concentration of 100uM.

#### Azide bead pulldown assay

Azide beads (Millipore Sigma, 900957) were resuspended by gentle flicking, until evenly mixed and washed twice with 10x volume of bead wash. This bead wash comprised of 0.1% BSA in DPBS solution containing 0.05% Tween-20 (Millipore Sigma, P1379), 50mM HEPES (ThermoFisher Scientific, J62688.AK), 400 mM Sodium Chloride (Millipore Sigma, S5150) and 1 mM EDTA (ThermoFisher Scientific, 15575020). Beads were spun down at 500g and resuspended with 1.5x added beadwash. 1 μL of 100 uM of each DBCO-NHS-PEG4 bound BOBcode was added to 20 μL of Azide beads in a PCR tube and then placed on a vortex machine for 30 minutes at 2500rpm. Each tube was then spun down at 500g for 30 seconds and the supernatant concentration was measured using a nanodrop. Percent oligo bound to bead wash was calculated by: [(Control) - (Test)] *100 / (Control). Variables are defined as: Control-concentration of BOBcode added to the tube along with azide agarose beads. Test-concentration of BOBcode in supernatant

#### Cell culture

Human HEK293T cells (ATCC #CRL-3216) and mouse RAW 264.7 cells (ATCC #TIB-71) were each cultured at 37°C and 5% CO2 in T-75 flasks with DMEM (Thermo #11965092) supplemented with 10% fetal bovine serum (VWR #89133-100) and 1% penicillin-streptomycin (Thermo #15070063). Cultures were maintained below 80% confluency. HEK cells were passaged with TrypLE (Thermo #12604013) using manufacturer specified protocols. RAW cells were passaged mechanically by cell scraping.

#### Compound & ASO treatment for multiplexed screen

HEK (30,000 cells/well) or RAW (60,000 cells/well) cells were plated in poly-D-Lysine (PDL)-coated (Thermo #A3890401) 96-well plates one day prior to azide labeling. After settling for four hours, cells were washed once with pre-warmed 1% BSA (R&D #5217) in DPBS (+Ca^2+^/Mg^2+^, Thermo #14040133) prior to transfection and/or compound treatment. Control cells were untreated. All treatment groups were completed in triplicate.

Where indicated, HEK cells were treated with 25 nM or 500 nM Risdiplam (MedChemExpress #HY-109101) in standard HEK medium for 24 hours. Where indicated, HEK cells were treated with 1 μg/mL or 50 μg/mL cycloheximide (Thermo #J66901.03) in serum-free DMEM for six hours.

Where indicated, HEK cells were transfected for 24 hours with 1 nM or 100 nM of a splice-switching ASO targeting *CADM1*. ASOs were prepared in a transfection mix using OptiMEM (Thermo #31985070) and mixed with an equal ratio of Lipofectamine 3000 (Thermo #L3000008) in OptiMEM, followed by incubation at room temperature for 15 minutes. The transfection mix was then added dropwise to plated cells in serum-free DMEM.

#### NAI-N_3_ RNA labeling

At the completion of each treatment, cells in all groups were washed once with pre-warmed 1% BSA in DPBS (+Ca^2+^/Mg^2+^) prior to NAI-N_3_ labeling. An azide-labeling mastermix was prepared comprised of 100 mM NAI-N_3_ (Enamine #EN300-247313) and 10 mM HEPES pH=7.0 (Thermo #<u>J62688.AK</u>) in pre-warmed 1% BSA in DPBS (+Ca^2+^/Mg^2+^). The mixture was prepared immediately prior to cell treatment and mixed well by pipetting to dissolve precipitates. The wash mix was aspirated from cells, and 50 μL of labeling mix was gently added to each well. Cells were incubated at 37°C for 10 minutes, then immediately washed three times in pre-warmed 1% BSA in DPBS (+Ca^2+^/Mg^2+^). After removing the final wash, 40 μL of Buffer RLT (Qiagen #79216) was added to each well and mixed by pipetting to lyse the cells. Lysates were stored at −80C.

#### BOBcode coupling via strained-promoted alkyne-azide cycloaddition (SPAAC)

Frozen cell lysates with azide-labeled RNA were thawed on ice, then transferred to PCR 8-strip tubes. 10 μL of a custom-prepared BOBcode stock (100 μM) was added to each sample, mixed 5 times by pipetting, vortexed briefly and spun down followed by an incubation for one minute at room temperature. Next, 40 μL of CleanNGS beads (Bulldog Bio #CNGS050) was added to each sample, mixed 5 times by pipetting, vortexed briefly and spun down. The SPAAC reaction mixture was incubated at 37°C for 30 minutes. Following completion, the reaction was quenched by mixing in 10 μL of a 1M sodium azide (Thermo #014314.09) solution, followed by pipetting 5 times to mix, a pulse vortex and spinning down. The quenching reaction was allowed to incubate for five minutes at room temperature, with periodic pulse vortexing every minute.

#### Sample pooling & RNA extraction

Individually BOBcoded labeled RNA samples were pooled following completion of the quenching reaction. From 24 pooled samples, a total volume of 2400 μL was pooled into a 15 mL conical tube. To prepare the mixture for column purification, the pooled sample was mixed with 4290 μL Buffer RLT, 60 μL nuclease-free water (Thermo #AM9939), then 3750 μL 100% ethanol (Sigma #E7023) to achieve a final ratio of RLT:water:ethanol of 350:100:250. The solution was mixed well to combine, then dispensed in 700 μL aliquots into individual RNeasy Mini Columns (Qiagen #74104).

RNA purification was completed following the Qiagen RNeasy Mini Kit manufacturer protocols. For the final elution, 30 μL of nuclease-free water was added to each column. Following a two minute incubation at room temperature, columns were spun down at 14,000g for one minute. The eluents of all columns were pooled into a single 1.5 mL eppendorf tube. Concentration and QC were determined by nanodrop and High Sensitivity RNA ScreenTape (Agilent #5067-5579) using the standard manufacturer protocol.

Due to carryover of BOBcodes apparent on TapeStation readings, a second round of column RNA purification was performed. 420 μL of the pooled RNA eluent from the first round was mixed with 1400 μL Buffer RLT and 1000 μL 100% ethanol, then dispensed in 700 μL aliquots into individual RNeasy Mini Columns. For the final elution, 60 μL of nuclease-free water was added to each column. Following a two minute incubation at room temperature, columns were spun down at 14,000g for one minute. The eluents of all columns were pooled into a single 1.5 mL eppendorf tube. Concentration and QC were determined by nanodrop and High Sensitivity RNA ScreenTape using the standard manufacturer protocol. Purified RNA was stored at −80C.

#### Selection of polyadenylated RNAs

Polyadenylated transcript enrichment from ∼18 µg of total RNA was performed using NEBNext Poly(A) mRNA Magnetic Isolation beads (NEB #E7490S) according to the manufacturer’s express protocol and with a maximum of 3 µg of total RNA input in each of 6 separate reactions. Following Poly(A) transcript selection, eluate fractions were combined and concentrated in 20 µL of water using an RNeasy Mini spin column (Qiagen #74104).

#### Reverse Transcription

cDNA was prepared from 700 ng of Poly(A) RNA using Maxima H Minus Reverse Transcriptase (Thermo #EP0751). The RNA sample was first mixed together with 2 µM R1-long-6N random hexamer, 1 µM dNTP (NEB #N0447S) and water up to 14 μL, then denatured at 72°C for three minutes. Denatured RNA was immediately placed on ice for 30 seconds, then combined with the provided 5X RT buffer, Maxima H Minus Reverse Transcriptase enzyme, and water up to 40 μL. The sample was mixed, then incubated at 25°C for 10 minutes, 42°C for 30 minutes, and 10 cycles of 50°C for 2 minutes followed by 42°C for 2 minutes. Enzyme inactivation was carried out by heating samples to 85°C for 5 minutes.

#### Exonuclease digestion

Excess primer from cDNA samples was removed using Exonuclease I digestion (NEB M0293L) by heating samples to 37°C for 30 minutes. Enzyme inactivation was carried out by heating samples to 80°C for 5 minutes. Following Exonuclease I digestion, cDNA samples were further purified by performing a 1.0X CleanNGS bead purification.

#### qPCR

The number of cycles used for amplification of libraries was determined by qPCR (ref: https://www.lexogen.com/blog/amplification-of-rna-seq-libraries-the-correct-pcr-cycle-number/). qPCR reactions were prepared using iTaq Universal SYBR Green supermix (Biorad #172-5124) according to the manufacturer’s directions. One-tenth of the cDNA library was used as input for each qPCR reaction. Reactions were run on a QuantStudio Real-Time PCR System 5 (Thermofisher). The cycle number at which 50% of the maximum fluorescence signal was achieved was determined. To account for the increased template concentration in the remaining nine-tenths of cDNA library that would be used for library amplification, three cycles were subtracted from this value. However, to ensure sufficient amplification, two additional cycles were added to the calculated number of cycles required for amplification.

#### Library amplification

cDNA was amplified using KAPA HiFi HotStart (Roche 07958897001).The thermalcycling protocol used for PCR amplification is the following: (1) 95°C for 3 minutes; (2) 3 cycles: 98°C for 20s, 68°C for 15s, 72°C for 1 minute; (3) 12 cycles: 98°C for 20s, 72°C for 1 minute; (4) 72°C for 2 minute.

#### Agarose gel electrophoresis

One-tenth of unpurified PCR product was subjected to gel electrophoresis using a 2% SYBR safe DNA stained E-gel (Invitrogen #A42135) and E-Gel Power Snap Electrophoresis System. PCR products were visualized using an iBright FL1500 Imaging System and size was assessed by running alongside a DNA ladder (NEB #N3231L).

#### Sample preparation and sequencing

The PCR amplified library was subjected to a double-sided bead cleanup using 0.6X and 0.8X bead ratios for right- and left-side size selections, respectively. Sequencing was performed by Novogene Corporation (Sacramento, CA) on a NovaSeq X Plus Series using a PE150 protocol.

#### High Sensitivity ScreenTape

An Agilent 4200 TapeStation was used to assess RNA and DNA samples using High Sensitivity RNA ScreenTape Assay and High Sensitivity D1000 ScreenTape Assay, respectively, according to manufacturer’s instructions.

#### Read processing and bobcode calling

The 24-plex BOBseq library (2 x 150 nt) was sequenced on one NovaSeq X Plus lane, yielding 182,207,880 read pairs. A bobcode was called at read-2 positions 1 to 7 against the 24 codes of the run, one substitution allowed, ties were rejected as ambiguous: 174,997,334 pairs (96.0%) received a code (168,298,599 exact, 6,698,735 with one mismatch) and 7,210,546 (4.0%) did not. The G-run was removed, poly(A) read-through was trimmed from the 3’ end of read 2, and pairs with a read 2 insert shorter than 30 nt after trimming were discarded, read 1 was left unmodified.

#### Alignment

All 24 samples were aligned with STAR 2.7.11b to a combined reference of the GRCh38 and GRCm39 primary assemblies with the Ensembl release 113 annotation, contigs prefixed by species. All alignments were reported (outSAMmultNmax unlimited, outFilterMultimapNmax 10000, unmapped reads retained) with score and matched-length thresholds of 0.33 of the pair length, at most 10 mismatches and 4% of the pair length, alignIntronMax 1,000,000 and local alignment ends.

#### Deduplication

PCR duplicates were removed through deduplicating by bobcode, contig, strand, 5’ position of read 2, fragment length and the UMI, exact match. Duplicate statistics were computed on the complete data before filtering. The 21 wells that passed input quality control retained 1.73 to 2.99 million molecules, the three risdiplam 500 nM wells (input failure) retained 0.37 to 0.54 million. These deduplicated molecules underlie the per-sample composition, the barcode-accuracy analysis, the benchmark uses its own molecule definition described below. [main bobseq fig, panel D]

#### Read classification and barcode accuracy

Molecules were classified by its primary alignment in one of the following categories: rRNA, mitochondrial (contig MT), ribosomal-protein mRNA, mRNA (exon of any other protein-coding gene), exonic of another biotype, intronic, intergenic. The species of a molecule was taken from all its alignments (human, mouse, or ambiguous when both). [Supplement fig], [main bobseq fig, panel D, E]

The per sample barcode accuracy was measured through a funnel of filtering steps on deduplicated molecules: mapped molecules with an assigned bobcode (L0); molecules aligning to one species only (L1); not rRNA or mitochondrial (L2); assigned to a protein-coding exon (L3); and carrying an exactly matching bobcode (L4). Accuracy at each level is the fraction of molecules belonging to the well’s expected species, with a 95% Wilson interval. [Main bobseq fig, Panel F]

#### Benchmark against published 3’ methods

Public datasets of benchmark methods were obtained for DRUG-seq (GSE176150, run SRR14730306: one 384-well U-2 OS plate, of which the 24 DMSO wells were used, well barcodes mapped to the published 384-barcode whitelist) and prime-seq (E-MTAB-10142: the eight HEK293T samples HEK_2, HEK_3, HEK_9, HEK_20, HEK_31, HEK_42, HEK_53 and HEK_64, runs ERR5375208 to ERR5375365, each 10,000 cells). For the BOBseq benchmark set,18 samples from the 24-plex run were used. The three risdiplam 500 nM wells were excluded as input failures, and the three mouse wells, which served only as species controls, were excluded from the human-only comparison.

Every method was processed through one path: the same reference and STAR multimapper settings (all alignments reported), one read table (well, UMI, contig, position, unique or multi-mapping, region class, gene, strand) and one unique molecule rule (sample/well, UMI, gene). The comparison is at native read length: DRUG-seq (52 nt) and prime-seq (50 nt) as single-end reads with score and matched-length thresholds of 0.66 of the read, BOBseq as the pipeline’s paired 2 x 150 nt alignments (thresholds 0.33 of the pair, the equivalent per mate). A filtered read is a uniquely mapped read (NH 1, MAPQ 255) assigned to an exon of a protein-coding gene on a human contig [Main Bobseq fig, Panel D]. For the distance-from-poly(A) coverage profile, a read-length-matched variant was also computed in which every method is one 50-nt single-end read per fragment (BOBseq read 2 truncated to 50 nt, all arms aligned with the 0.66 thresholds) [Supplement fig, distance from polyA 50 nt fig].

Depth matching used rarefaction: the filtered reads of every well were subsampled without replacement to 50,000, 100,000, 250,000, 500,000, 1 million and 2 million reads, and molecules and genes (ribosomal-protein genes not counted) were determined at each depth; curves show the mean over the wells of a method with its 95% t-interval [Supplement fig, stacked plot]. Splice junctions were extracted per sample from the deduplicated, uniquely mapped alignments with regtools 1.0.0 (junctions extract, minimum anchor 4 nt, intron length 1 to 10^^9^ nt, strand from the XS tag), a read pair spanning a junction counted once. A junction was counted when supported by at least three molecules. Splicing curves pooled the wells of each method, reaching 4,405, 22,925 and 143,452 junctions with at least three molecules at full depth. A junction was called annotated when its intron coordinates matched an intron of the Ensembl 113 annotation (528,735 human introns). [Main Benchmark fig, Panel D] The intronic fraction per well is the intronic class of the classification above as a percentage of mapped reads, multimappers included. [Main Benchmark fig, Panel C]

#### Coverage along transcripts

Coverage was analyzed on one canonical transcript per protein-coding gene. The poly(A)-anchored profile is computed per method from the reads of its benchmark wells pooled: genes with a canonical transcript of at least 1,000 nt, ribosomal-protein genes excluded, and at least 200 unique reads in every method (2,070 genes). The per-base coverage of each gene was divided by the gene’s mean coverage. [Main Benchmark fig, Panel A] The 5’ to 3’ balance was computed per well with Picard CollectRnaSeqMetrics 3.5.0 (STRAND_SPECIFICITY NONE, MINIMUM_LENGTH 1000) against a refFlat of protein-coding Ensembl 113 transcripts without ribosomal-protein and mitochondrial genes, as twice the coverage-weighted mean position over the 101 bins. [Main Benchmark fig, Panel B] The intronic fraction is per well as described above. [Main Benchmark fig, Panel C] The genome-browser views are pooled and depth-matched at the method level: the per-sample human alignments of the benchmark wells were pooled per method and subsampled to 8 million uniquely mapped fragments, selected by a 64-bit hash of sample and read name. [Main Benchmark fig, Panel E, F]

#### Software and data availability

STAR 2.7.11b, regtools 1.0.0, samtools 1.24, Picard 3.5.0, deepTools 3.5.6, Python 3.12 with pysam, numpy, scipy and matplotlib. Sequencing data are available at NCBI SRA under BioProject PRJNA1532586 (SRA study SRP738601): the 24 per-sample read pairs of the 24-plex library and the 195 nanopore datasets of the bobcode screen and the random-priming comparison (runs SRR40770546 to SRR40770764). The gene by sample count matrix is provided in the repository and as a supplementary table.

### Comparison to other HEK293 datasets

Principal-component analysis was performed on 29 untreated HEK293 control libraries (BOBseq, n = 3; prime-seq, n = 8; BRB-seq, n = 8; TruSeq, n = 10, spanning three GEO series). Transcripts were quantified with Salmon and summarized to gene level with tximport over the project mRNA gene set—Ensembl 113 primary-assembly protein-coding genes with the 13 mitochondrially encoded genes and 169 ribosomal-protein genes (^(RPL|RPS|MRPL|MRPS), excluding RPS6K) removed and haplotype copies collapsed (∼19,934 genes). Expression was expressed as per-million values within this gene set, using length-normalized TPM for the near-full-length methods (BOBseq and TruSeq) and CPM for the 3′-tag methods (prime-seq and BRB-seq); to remove the ∼100-fold depth range, every sample was first down-sampled (“thinned”) to the smallest mRNA library (325,111 mRNA reads) before per-million normalization. Values were log2(x + 1)-transformed, genes with mean expression ≥ 1 were retained, and the 5,000 most variable of these (by variance of log2 expression) were selected. Each gene was mean-centered and scaled to unit variance, and PCA was computed with scikit-learn (full singular-value decomposition, `random_state = 0`); sample scores were plotted for PC1 versus PC2 (29% and 17% of variance, respectively), colored by library method, with the ten TruSeq libraries additionally distinguished by a separate marker for each source laboratory (GEO series). Analyses used Salmon and tximport (v1.40.0; R 4.6.0/Bioconductor 3.23) and Python 3.13.9 (scikit-learn 1.9.0, numpy 2.5.3, pandas 2.3.3, matplotlib 3.10.9).

Differential expression between library methods was quantified with DESeq2 (v1.52.0) on the depth-thinned gene counts using the design `∼ condition`, Wald tests, and apeglm-shrunk log2 fold changes, retaining genes with at least 10 counts; per-gene, per-method length offsets were applied to the BOBseq and TruSeq samples so that all methods were compared on length-normalized expression. Three contrasts were computed against TruSeq as the common reference (BOBseq vs TruSeq, prime-seq vs TruSeq, and BRB-seq vs TruSeq; log2FC > 0 denotes higher expression than TruSeq), each restricted to a capped common gene set of ∼7,500 genes—defined as the intersection of the most highly expressed genes per method (accounting for ∼90–95% of each sample’s counts)—with DESeq2 re-fitted on this set so that the same genes were tested in all three contrasts. Genes with a Benjamini–Hochberg-adjusted P < 0.001 were called differentially expressed. Three three-set Venn diagrams were then generated from these lists: one of all differentially expressed genes (either direction) and two split by direction (genes higher, and genes lower, than TruSeq). Region membership was computed as the exact set intersection minus the union of the remaining circles (exclusive counts), and diagrams were drawn with a custom, non-area-proportional three-set Venn in matplotlib (v3.10.9) in which each region is annotated with its exclusive gene count and each circle with its total; for the all-genes diagram, each shared region was additionally annotated with the number of genes showing concordant versus discordant fold-change direction across the methods of that region. Genes tested were those with a non-missing adjusted P value in all three contrasts, and region membership, direction concordance, and gene identities were exported to a supplementary table. Analyses used R 4.6.0/Bioconductor 3.23 (DESeq2 1.52.0, apeglm 1.34.0, tximport 1.40.0) and Python 3.13.9.

### Sample to sample variance calculation

#### Datasets

Replicate transcriptomes were compared for three bulk RNA-seq library-preparation methods. BOBseq libraries were generated in this study (HEK cells; a 24-plex experiment comprising untreated controls and drug-treatment conditions, each in triplicate; full-length, randomly-primed, paired-end libraries with a 6-nt unique molecular identifier [UMI]). prime-seq data were obtained from ArrayExpress accession E-MTAB-10142 (Janjic et al., 2022)¹; we used the eight untreated HEK293T samples of a single lysis condition (“Incubation + Proteinase K”, 10,000 cells per lysate), which are 3′-end poly-dT libraries (50-nt single-end, 10-nt UMI). DRUG-seq data were obtained from Gene Expression Omnibus accession GSE176150 (Li et al., 2022)²; we used the 24 DMSO vehicle-control wells (24 h) of one U-2 OS plate (VH02001704_S4), a 3′-end tag method with UMIs (Ye et al., 2018)³ originally describes the assay. One BOBseq risdiplam-500 triplicate that failed input QC was excluded; the BOBseq Hek_control_2 well (a T-depleted UMI oligo) was retained as it was otherwise a normal control.

Uniform re-processing and quantification

To remove processing as a confounder, all datasets were re-processed from raw reads with an identical pipeline. Reads were aligned to the human genome (GRCh38, Ensembl release 113) with STAR v2.7.11b⁴. For the UMI-containing libraries (BOBseq, prime-seq, DRUG-seq), aligned reads were deduplicated to one read (or read pair) per unique combination of mapping position, strand, and UMI. Uniquely mapped reads (MAPQ 255) were assigned to genes with featureCounts⁵ against the Ensembl 113 gene annotation, yielding an integer gene-by-sample count matrix.

#### mRNA gene set

To compare methods on the messenger transcriptome and to exclude gene classes captured very differently across chemistries, an “mRNA” gene set was defined as protein-coding genes (Ensembl biotype protein_coding) after removing (i) cytoplasmic and mitochondrial ribosomal-protein genes (symbols matching RPL, RPS, MRPL, or MRPS, excluding RPS6K kinases) and (ii) all mitochondrially-encoded genes (chromosome MT), giving 19,934 genes. This exclusion was motivated by strong, method-specific composition differences in the raw libraries (e.g., ∼19% of prime-seq and ∼2% of DRUG-seq gene-assigned reads were mitochondrial, versus <1% for BOBseq; ∼29% of DRUG-seq reads were ribosomal-protein), which would otherwise dominate correlations. Counts within the mRNA set were converted to counts per million (CPM) re-normalized within that set, and log-transformed as log₂(CPM + 1).

#### Depth normalization

Analyses were performed either at native sequencing depth (no downsampling) or after depth-matching. For depth-matching, each sample’s mRNA gene-count vector was thinned to a common target by multinomial subsampling without replacement (variance-preserving, fixed random seed). Depth targets were chosen as the minimum mRNA-assigned depth of a defined reference set — the smallest prime-seq library (296,992 counts), the smallest of the eight highest-depth DRUG-seq wells (141,053 counts), or the smallest library across the full sample set (48,837 counts) — as specified per analysis.

#### Gene inclusion and reproducibility metric

For each pair of replicate samples, genes were included either by a detection threshold (≥1, ≥3, or ≥5 counts in both samples of the pair) or by a rank threshold (the N most highly expressed genes for that pair, ranked by the mean of the two samples’ CPM, with no count cutoff). When a common rank threshold across a shallow set was used, N was capped at the largest value for which every pair had that many genes detected in both samples, to avoid inflating correlations with dropout (genes present in only one replicate). Replicate-to-replicate reproducibility was quantified as the Pearson correlation coefficient of log₂(CPM + 1) over the included genes. For BOBseq, correlations were computed only between replicates within the same experimental condition (the triplicate of each of six conditions) and pooled across conditions; for prime-seq and DRUG-seq, which are each a single condition, all within-method replicate pairs were used.

#### Statistics

Because replicate pairs are not independent (each sample contributes to multiple pairwise comparisons), significance was assessed on a per-sample summary rather than on individual pairs: for each sample we computed its mean correlation to the other replicates of its group, and compared these per-sample distributions between methods with a two-sided Mann–Whitney U test. Interquartile range of the per-pair correlations was reported as a measure of within-method consistency. Analyses used Python 3 with NumPy, SciPy⁶ (scipy.stats.mannwhitneyu), and Matplotlib.

### 24 plex gene expression comparison

#### CHX dose response on gene expression

Gene-expression changes across the cycloheximide dose series (control, 1 µg/mL, and 50 µg/mL; nine wells, three per dose) were quantified from BOBseq gene counts. Reads were aligned with STAR 2.7.11b to a combined GRCh38.113 + GRCm39 index; human primary alignments were deduplicated on UMI plus position (pysam 0.24.0) and filtered to uniquely mapped reads (`samtools view -q 255`, NH = 1), and UMI-deduplicated fragments were counted per gene with featureCounts 2.1.1 (`-t exon -g gene_id -s 1 -Q 255 --primary - p --countReadPairs`) against the Ensembl 113 annotation. Differential expression was tested with DESeq2 1.52.0: genes with at least 10 counts summed over the nine wells were retained, and the design `∼ dose` encoded dose as a numeric rank (0/1/2 for control/1/50 µg/mL), so the model fits a monotone dose trend. Significance was assessed by the Wald test on the dose-slope coefficient using DESeq2 defaults (median-of-ratios size factors, dispersion trend and shrinkage, Cook’s-distance outlier handling, and independent filtering at α = 0.1), and P values were Benjamini–Hochberg (BH) adjusted. The plotted effect size (“CHX dose-trend log2FC”) is twice the apeglm-shrunk slope, i.e. the modeled top-dose-versus-control log2 fold change. In the volcano plot, each point is one tested gene, with the dose-trend log2 fold change on the x-axis and −log10 BH-adjusted P on the y-axis; differentially expressed genes (BH-adjusted P < 0.001, no fold-change threshold) are colored by direction, a dotted guide marks the significance cutoff, and the labeled genes are the top hits by adjusted P. Analyses used R 4.6.0/Bioconductor 3.23 (DESeq2 1.52.0, apeglm 1.34.0) with upstream STAR 2.7.11b, samtools 1.24, subread/featureCounts 2.1.1, and pysam 0.24.0, and Python 3.13.9 (matplotlib 3.10.9).

#### CHX dose response on PSI values

Differential splicing across the cycloheximide dose series (control, 1 µg/mL, and 50 µg/mL; nine wells, three per dose) was quantified from BOBseq junction counts. Reads were aligned with STAR 2.7.11b to a combined GRCh38.113 + GRCm39 index, and human primary alignments were filtered to uniquely mapped reads (`samtools view -q 255`, NH = 1); split-read junctions were extracted from the CIGAR strings (regtools 1.0.0 semantics, minimum anchor 4 nt, counting distinct read pairs) and assigned to local splicing variations (LSVs; junctions sharing a source or target anchor) in the atlas v1 catalogue (GRCh38/Ensembl 113). For each LSV side, per-well percent-spliced-in (PSI) was computed as the junction fragment count divided by the summed counts of the detected members of that side, with the fragment (per read pair) as the counting unit; UMI-deduplicated ΔPSI was reported alongside as a check. Within the contrast, LSV sides were preselected in a condition-blind manner (≥ 100 fragments on average, detected in every well, and PSI variance across wells ≥ 0.005). Each retained event’s per-well PSI was regressed on dose rank (0/1/2 for control/1/50 µg/mL) by ordinary least squares, and the slope was tested with a two-sided t-test (a monotone dose-trend test that leverages the intermediate dose), with Benjamini–Hochberg correction applied within the contrast. The effect size, ΔPSI, was defined as PSI(top dose) − PSI(control) from the pooled fragment counts of each condition’s wells. In the volcano plot, each point is one LSV-side/junction, with ΔPSI (top dose − control) on the x-axis and −log10 BH-adjusted P on the y-axis; events passing the significance and effect-size thresholds (BH-adjusted P < 0.1 and |ΔPSI| ≥ 0.1) are highlighted and guides are drawn at ΔPSI = ±0.1 and P = 0.1. Analyses used Python 3.13.9 (numpy 2.5.3, scipy 1.17.1, matplotlib 3.10.9), with upstream STAR 2.7.11b, samtools 1.24, regtools 1.0.0 (semantics), and pysam 0.24.0.

#### SMN2 junction calculation

SMN2 exon-7 splicing was quantified paralog-aware from junction reads (extraction, c.840/c.*239 paralog assignment, and UMI deduplication as above). For the junction-composition panel, deduplicated inclusion (SMN2 ex6>7, SMN2 ex7>8) and skipping (ex6>8) reads were pooled across the three wells of each condition and expressed as proportions of their sum; because exon-7–skipping reads carry no paralog-diagnostic base, the skip class is the shared SMN1/SMN2 pool while the inclusion classes are SMN2-specific. For the PSI panel, per-well SMN2 exon-7 PSI was computed as SMN2 inclusion / (SMN2 inclusion + shared skip) reads, and risdiplam (Ris 25, n = 3) was compared with all other treated and control wells (n = 15; the QC-failed Ris 500 wells excluded) by a two-sided Mann–Whitney U test; points are per-well PSI and the crossbar is [mean ± SEM — confirm].

### Incubation time estimates

Protocol steps for each method were compiled from the published protocols and annotated with their stated reaction/incubation times (QC and final library cleanup excluded). Steps were grouped by type (reverse transcription, PCR, ExoI, purification, other) and classified as performed per sample or on the pooled library according to each method’s barcoding chemistry; the per-sample total (bar rail) reflects work that scales with sample number, whereas pooled steps are performed once regardless of multiplexing. Incubation times only are presented as hands on labor time is not easily estimated as presented in published methods.

---

**Supplemental Figure 1.**
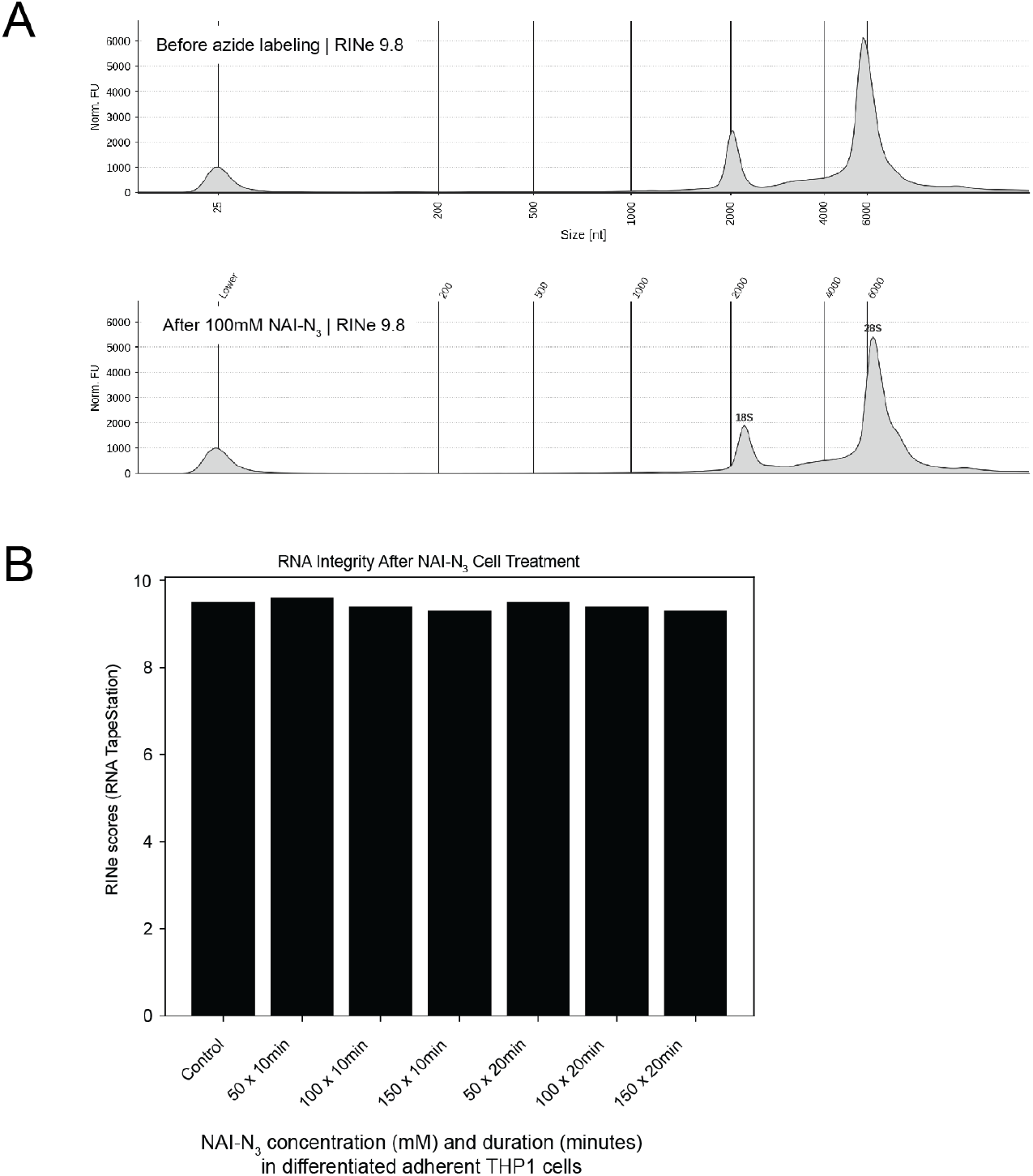
NAI-N_3_ treatment of RNA does not affect RINe scores. (A) RNA TapeStation electrophoresis plot profiles of total RNA before and after NAI-N_3_ labeling. (B) Differentiated adherent THP-1 cells were treated with different concentrations and durations of NAI-N3 before purifying RNA and calculating RINe scores with TapeStation.

**Supplemental Figure 2.**
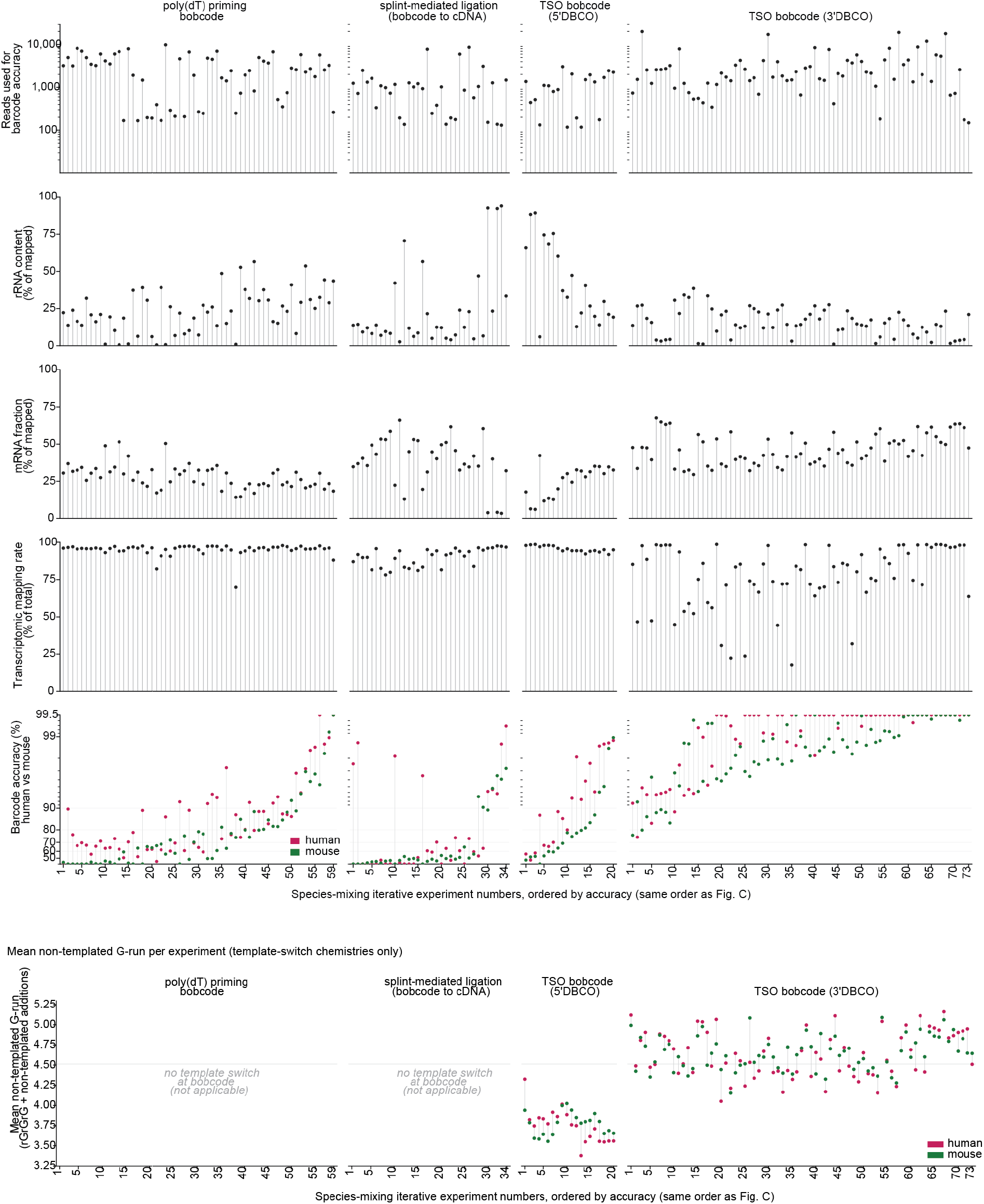
Intramolecular bobcode sequence transfer metrics for each individual experiment. Sequencing experiment metrics across each individual iterative experiment.

**Supplemental Figure 3.**
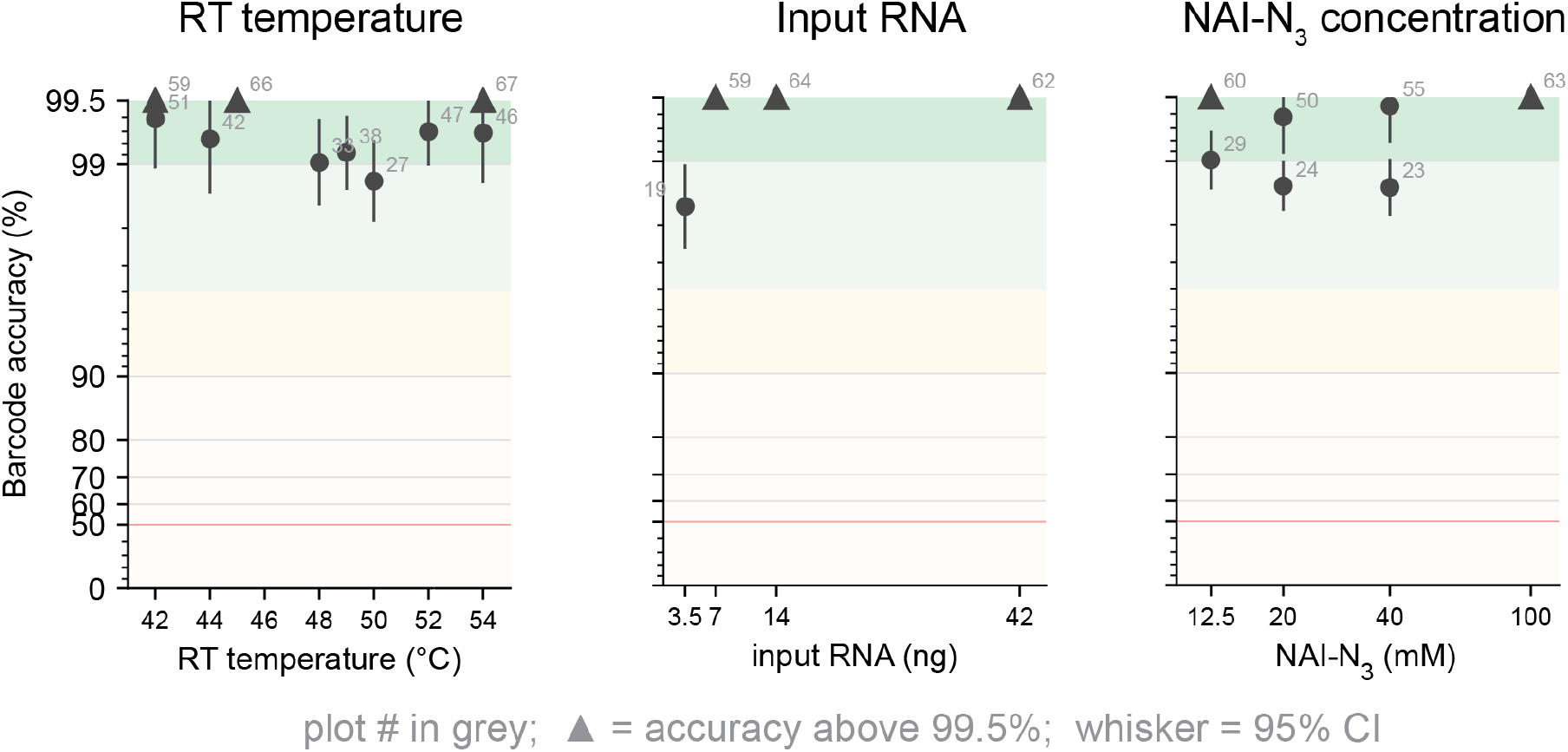
Subset of experiments from the bobcode screen. Rearranged subset of the bobcode accuracy results as in the main figure.

**Supplemental Figure 4.**
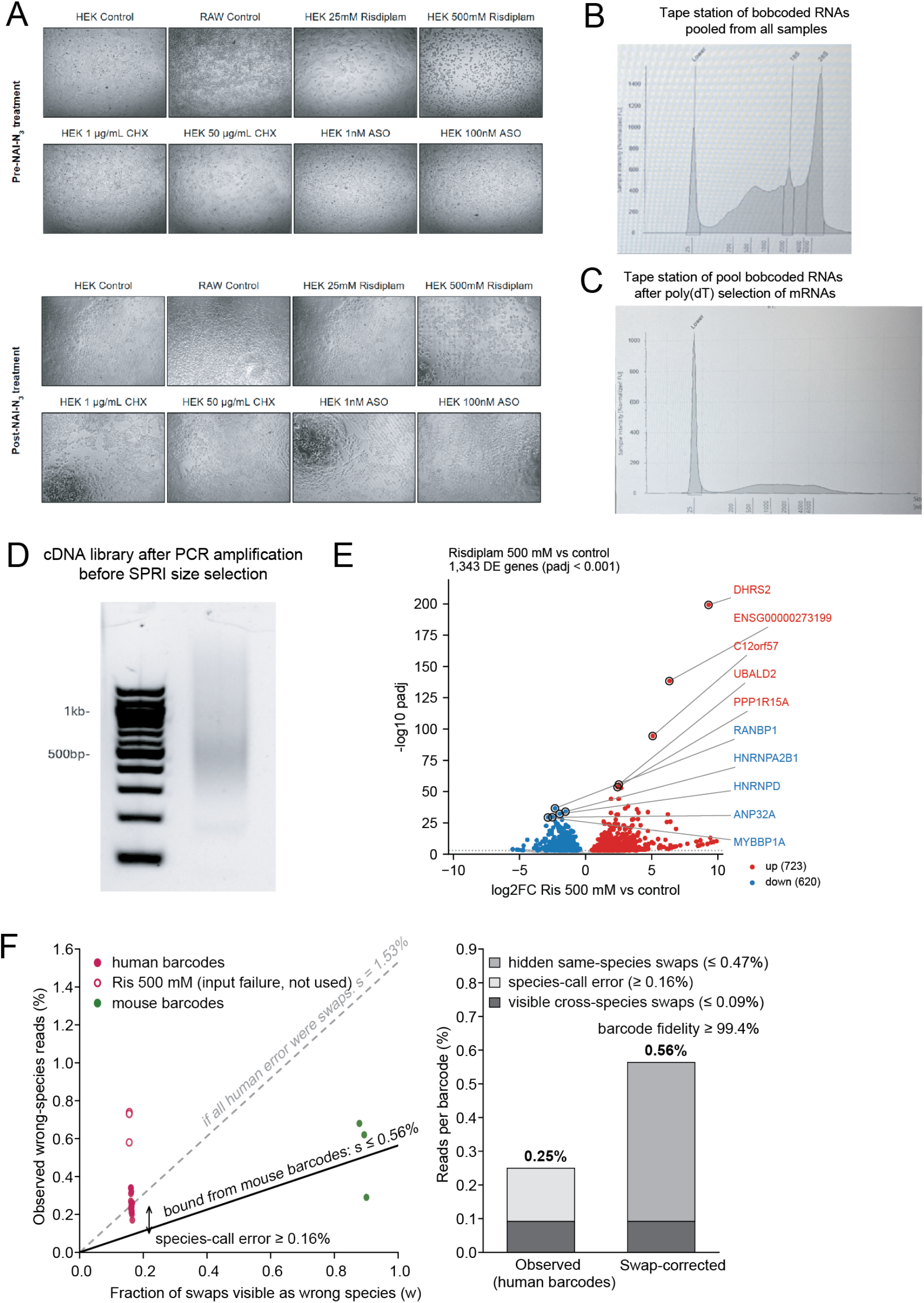
24plex BOB-seq drug screen. (A) Microscopy images of cells before and after NAI-N_3_ treatment. (B) TapeStation profile of RNA after pooling bobcoded RNA samples and multiplexed purification. Higher than usual peaks between ∼100-1500bp are hypothesized to be bobcode conjugation artifacts. (C) After Poly(dT) pulldown of polyadenylated mRNAs, rRNA peaks and bobcode artifacts are no longer present. (D) After PCR but before any SPRI size selection, we examined the distribution of the cDNA library. We observe an even distribution with peak at ∼500bp and tail. Samples were double-sided SPRI selected after this step prior to Illumina sequencing. (E) High dose risdiplam gene expression changes compared to controls. (F) Corrected barcode swapping rates from human and mouse samples identifying fidelity of >99.4% after correction.

**Supplemental Figure 5.**
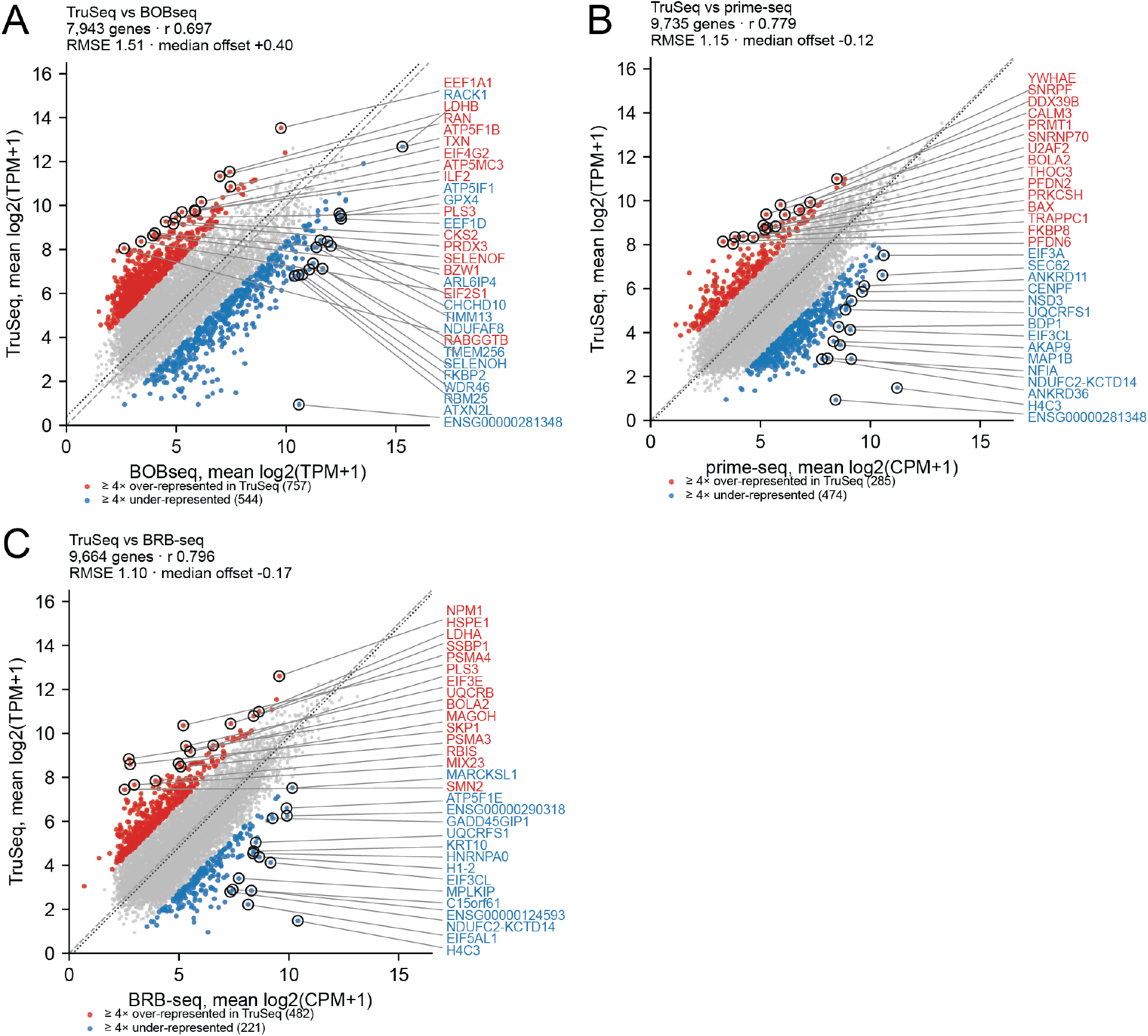
Comparing BOB-seq with other multiplexed RNA sequencing platforms from HEK293 cells. (A-C) Differential expression of TruSeq data with BOBseq (A), prime-seq (B), and BRB-seq (C).

**Supplemental Figure 6.**
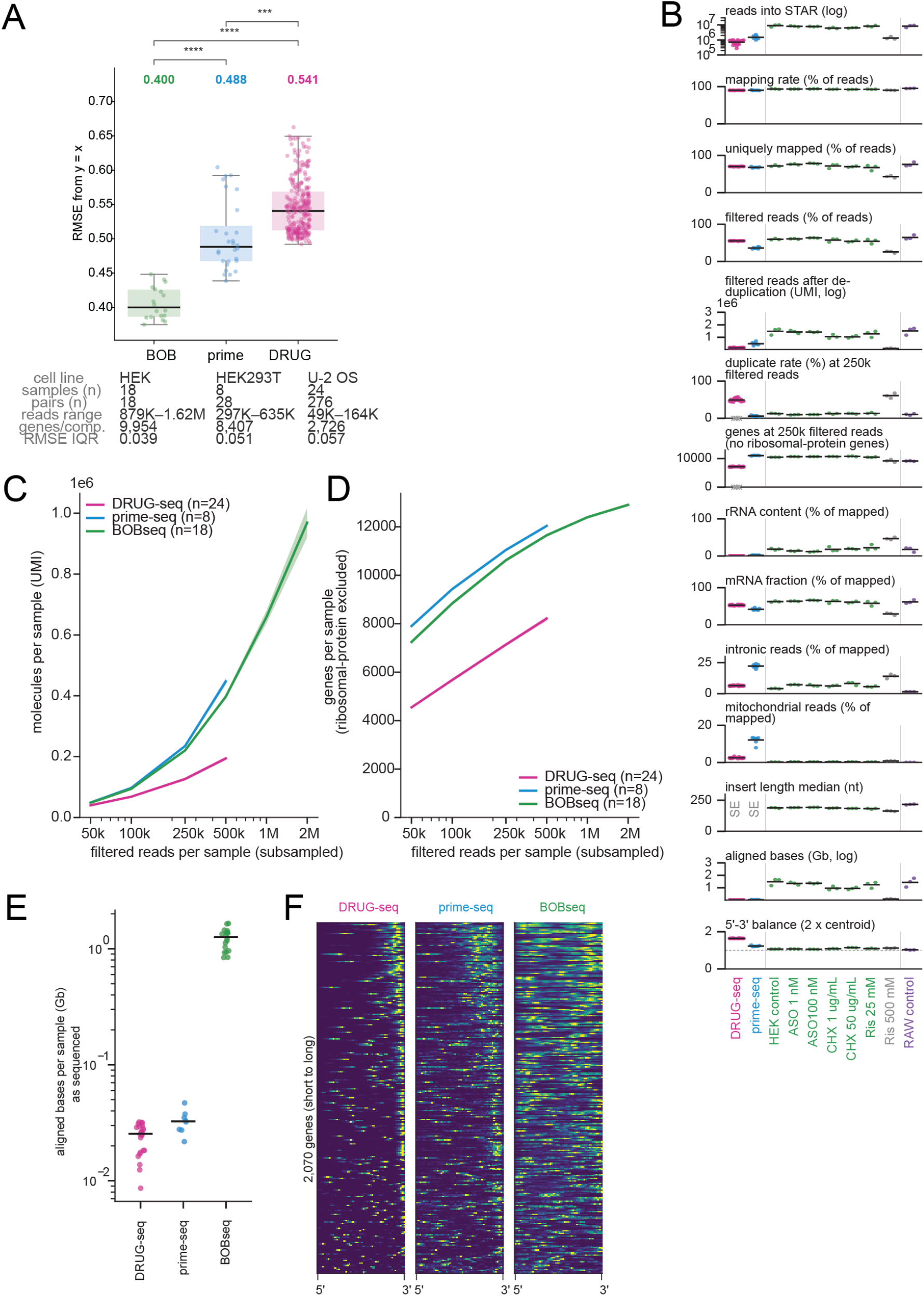
Comparing BOB-seq versus other multiplexed transcriptomic drug screening methods. (A) RMSE variance of gene expression within methods. (B) Sequencing alignment metrics across platforms and conditions. (C) Unique molecules per sample versus filtered reads per sample with 95% CI. (D) Genes detected per sample versus filtered reads. (E) Total number of aligned bases is higher in BOB-seq due to use of PE150 sequencing compared to shorter single ended sequencing with other methods. (F) Plotting gene coverage of the top 2070 genes ranked from short to long, demonstrating the 3’ bias in DRUG-seq and prime-seq compared to BOB-seq.

**Supplemental Figure 7.**
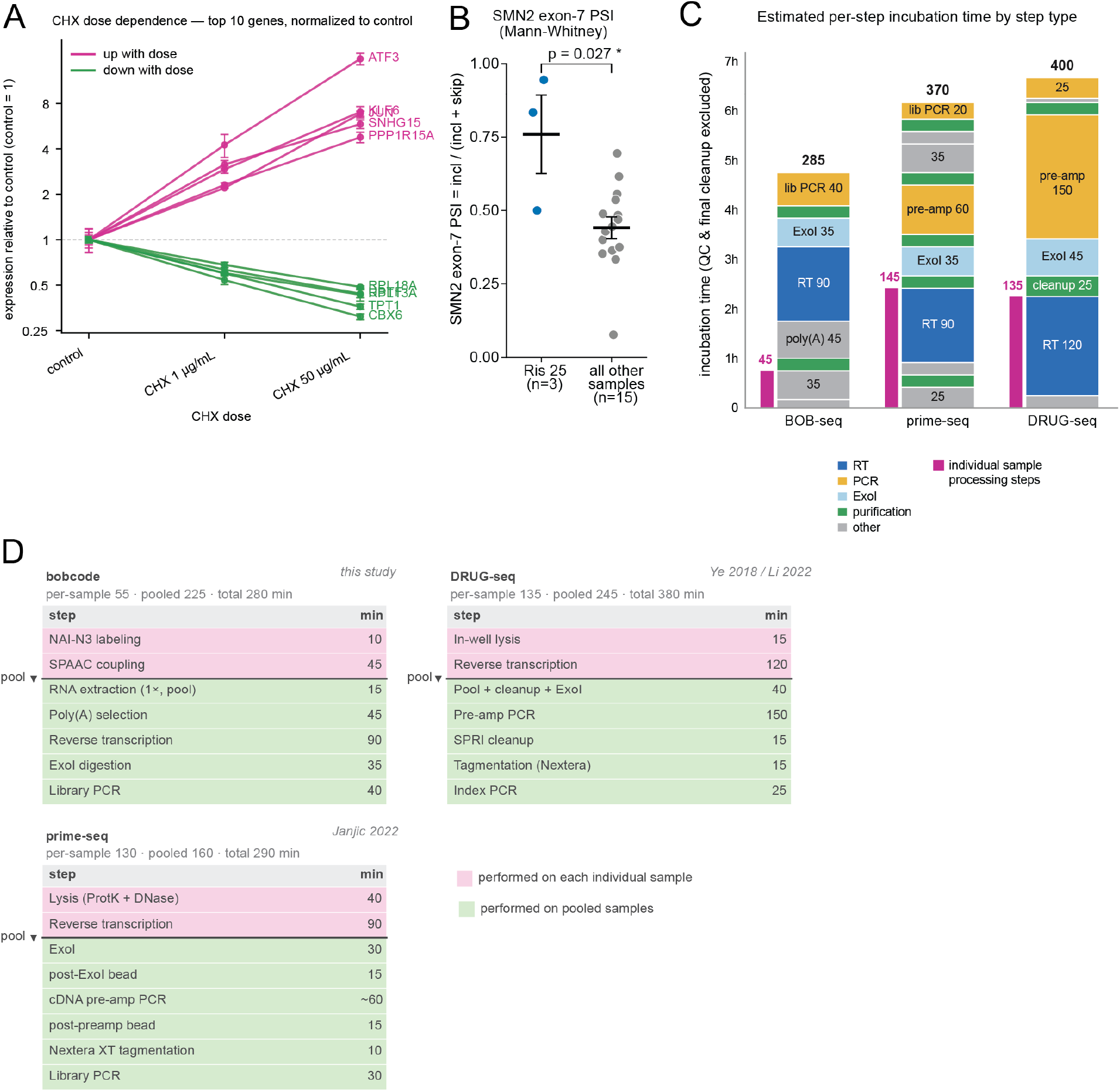
BOBseq expression, splicing, and workflow characteristics. (A) CHX dose-dependent gene expression of top affected genes in each condition. (B) Ris25 samples exhibit higher exon 7 inclusion compared to all other samples. (C-D) Estimated incubation time differences (y axis= hours, number in the plot in minutes) between BOB-seq, prime-seq, and DRUG-seq.

